# Tissue-resident CD4^+^ T helper cells assist protective respiratory mucosal B and CD8^+^ T cell memory responses

**DOI:** 10.1101/2020.02.28.970400

**Authors:** Young Min Son, In Su Cheon, Yue Wu, Chaofan Li, Zheng Wang, Yao Chen, Yoshimasa Takahashi, Alexander L. Dent, Mark H Kaplan, Yang-Xin Fu, Justin J. Taylor, Weiguo Cui, Jie Sun

## Abstract

The roles of CD4^+^ T helper cells (T_H_) in shaping localized memory B and CD8^+^ T cell immunity in the mucosal tissues are largely unexplored. Here, we report that lung T_H_ cells provide local assistance for the optimal development of tissue-resident memory B (B_RM_) and CD8^+^ T (T_RM_) cells following the resolution of primary influenza virus infection. We identify a population of tissue-resident CD4^+^ T_H_ (*aka* T_RH_) cells that co-exhibit follicular T helper (T_FH_) and T_RM_ cell features and mediate local help of CD4^+^ T cells to B and CD8^+^ T cells. Optimal T_RH_ cell formation requires lung B cells and transcription factors involved in T_FH_ or T_RM_ development. Further, we show that T_RH_ cells deliver local help to B and CD8 T cells through CD40L and IL-21-dependent mechanisms. Our data have uncovered a new tissue-resident T_H_ cell population that is specialized in assisting the development of mucosal protective B and CD8^+^ T cell responses *in situ*.

## Introduction

The long-term protection against pathogen reinfection is mediated by long-lived plasma cells, memory B cells (B_MEM_) and/or memory T (T_MEM_) cells. In addition to circulating B_MEM_ and T_MEM_ cells, tissue-resident memory B (B_RM_) and T (T_RM_) cells that reside in the mucosal sites have recently been identified and characterized ^1, 2, 3, 4, 5^. B_RM_ and T_RM_ cells are able to mount rapid recall responses *in situ* against invading pathogens before pathogen dissemination, and therefore are thought to provide immediate and superior protection against secondary infections ^1, 6, 7, 8^. However, the mechanisms underlying the development and persistence of robust B_RM_ and T_RM_ cell responses in the respiratory tract are largely undefined. Furthermore, we do not know whether there are cellular and molecular pathways that can be targeted to simultaneously promote both B_RM_ and T_RM_ responses for maximal protection against pathogen reinfections.

Influenza viruses remain a leading cause of respiratory tract infections despite progress in antiviral therapies. Each year, influenza virus infects 5–10% of adults and 20–30% of children worldwide ^9, 10^, resulting up to 35.6 million illnesses and 56,000 deaths annually in the U.S. since 2010 ^11^. Influenza virus infection induces potent development of protective B_RM_ and CD8^+^ T_RM_ responses in the respiratory mucosal tissue ^5, 12, 13^. Compared to B_MEM_ cells in the secondary lymphoid organs, lung B_RM_ have enhanced percentages of cells poised for cross-reactive memory repertoires, potentially due to the local supply of B_RM_ precursors by persistent germinal center (GC) responses in the inducible bronchus-associated lymphoid tissues (iBALT) following primary influenza infection ^14^. Therefore, influenza-specific B_RM_ cells are thought to provide better cross-protection at the site of infection compared to their counterparts in the secondary lymphoid organs ^1, 14^. Additionally, lung B_RM_ cells, but not systemic B_MEM_ cells, contributed to early plasmablast responses following influenza re-infection, which are potentially important in restricting early viral dissemination ^1, 5, 14^. T_MEM_ responses against conserved influenza internal epitopes confer cross-reactive protection against influenza viruses that escape neutralizing antibody (Ab) responses ^15, 16, 17, 18, 19, 20^. In animal models, influenza-specific lung CD8^+^ T_RM_ cells can rapidly respond to heterologous influenza virus reinfection before it can replicate to high titers ^21, 22^. Influenza-specific CD8^+^ T_RM_ cells have also been detected in human lungs and are thought to be important for the protection against severe influenza-associated diseases ^23, 24^. However, compared to other tissues, lung protective CD8^+^ T_RM_ responses are short-lived in nature ^25, 26^. Thus, understanding the cellular and molecular mechanisms regulating the development and/or maintenance of lung protective B_RM_ and/or CD8^+^ T_RM_ responses may aid the design of future influenza vaccines.

CD4^+^ T helper (T_H_) cells are important in anti-influenza immune responses by providing essential “help” for the development of effector and memory CD8^+^ T and germinal center B (B_GC_) cell responses ^15^. CD4^+^ T cells are not required for the generation of effector CD8^+^ T cells during primary influenza infection ^27^, but are needed for the production of IL-10 by influenza-specific effector CD8 T cells ^28^. CD4^+^ T cell help, particularly during the priming stage, is vital for the development of circulating T_MEM_ and CD103^+^ T_RM_ following primary influenza infection ^29, 30^. Similarly, CD4^+^ T cell help, mediated mainly through follicular helper T cells (T_FH_) in the secondary lymphoid organs, is required for assisting B cells to form GC and the production of high affinity antibodies during influenza infection ^31, 32, 33^. However, whether CD4^+^ T cells can assist local B and CD8^+^ T cell memory responses at the mucosal tissue following the resolution of primary infection is unknown. Several recent studies have also identified a population of PD-1^hi^ “T_FH_-like” cells in the peripheral tissues during autoimmunity ^34, 35, 36^. However, the developmental cues and the precise physiological functions of these cells remain largely elusive. During influenza infection, the existence of a lung “T_FH_-like” cell population that can potentially sustain lung B_GC_ responses has been recently demonstrated ^35, 37, 38^, but is still controversial ^5, 36^.

Here, we have identified a population of lung CD4^+^ helper T (T_H_) cells developed after influenza viral clearance, co-exhibiting T_FH_ and T_RM_ features. Based on their gene expression, sessile features and functional properties, we termed these cells tissue-resident T helper cells (T_RH_). Importantly, T_RH_ cells provide local help for the generation of strong B_GC_ and B_RM_ responses, as well as a CD8^+^ T_RM_ population that is critical for the protection against heterologous influenza virus infection ^39, 40^. We further identified the molecular cues mediating T_RH_ cell help to B and CD8^+^ T cells. Our findings reveal a previously unidentified tissue-resident T_H_ cell population that is important in assisting local development of protective memory B and CD8^+^ T cell responses in the respiratory mucosal tissue.

## Results

### Lung CD4^+^ T cells provide “late” help for the formation of B_GC_, B_RM_ and CD8^+^ T_RM_ responses *in situ*

We sought to determine whether CD4^+^ T cell help can assist local B and CD8^+^ T cell memory responses at the mucosal tissue following the clearance of primary infection. To this end, we utilized a mouse model of influenza A virus PR8/34 (PR8) infection, in which viral clearance occurs within 10 days post infection (d.p.i.) ^41, 42, 43^. We infected WT mice with PR8 virus and depleted CD4^+^ T cells with α-CD4 (GK1.5, 250 μg/mouse/weekly) injection starting at 14 d.p.i. (Fig. 1a). CD4^+^ T cells were largely depleted in the spleen and the lung as confirmed by flow cytometry (Extended Fig. 1a). At 42 d.p.i., we analyzed lung tissue B and T cell responses through intravenous (i.v.) injection of α-CD45 5 min before mouse sacrifice as we and others reported ^40, 44, 45^. In this setting, CD45_i.v._^-^ cells were within lung tissue, while CD45_i.v._^+^ cells were in lung blood vessels. CD4^+^ T cell depletion disrupted the formation of iBALT, which contained B cell and CD4^+^ T cell aggregates ^37, 46, 47^ (Fig. 1b and Extended Fig. 1b). CD4^+^ T cell depletion also abrogated lung B_GC_ responses (Extended Fig. 1c). We then examined influenza Hemagglutinin-specific B (HA-B) cell responses in the lungs and spleens following CD4^+^ T cell depletion at 42 d.p.i. CD4^+^ T cell depletion did not affect lung circulating (CD45_i.v._^+^) nor splenic HA-specific B cell responses, but drastically diminished total and HA-specific B cells in the lung tissue at the memory stage (Fig. 1c and Extended Fig. 1d, e).

**Fig. 1.**
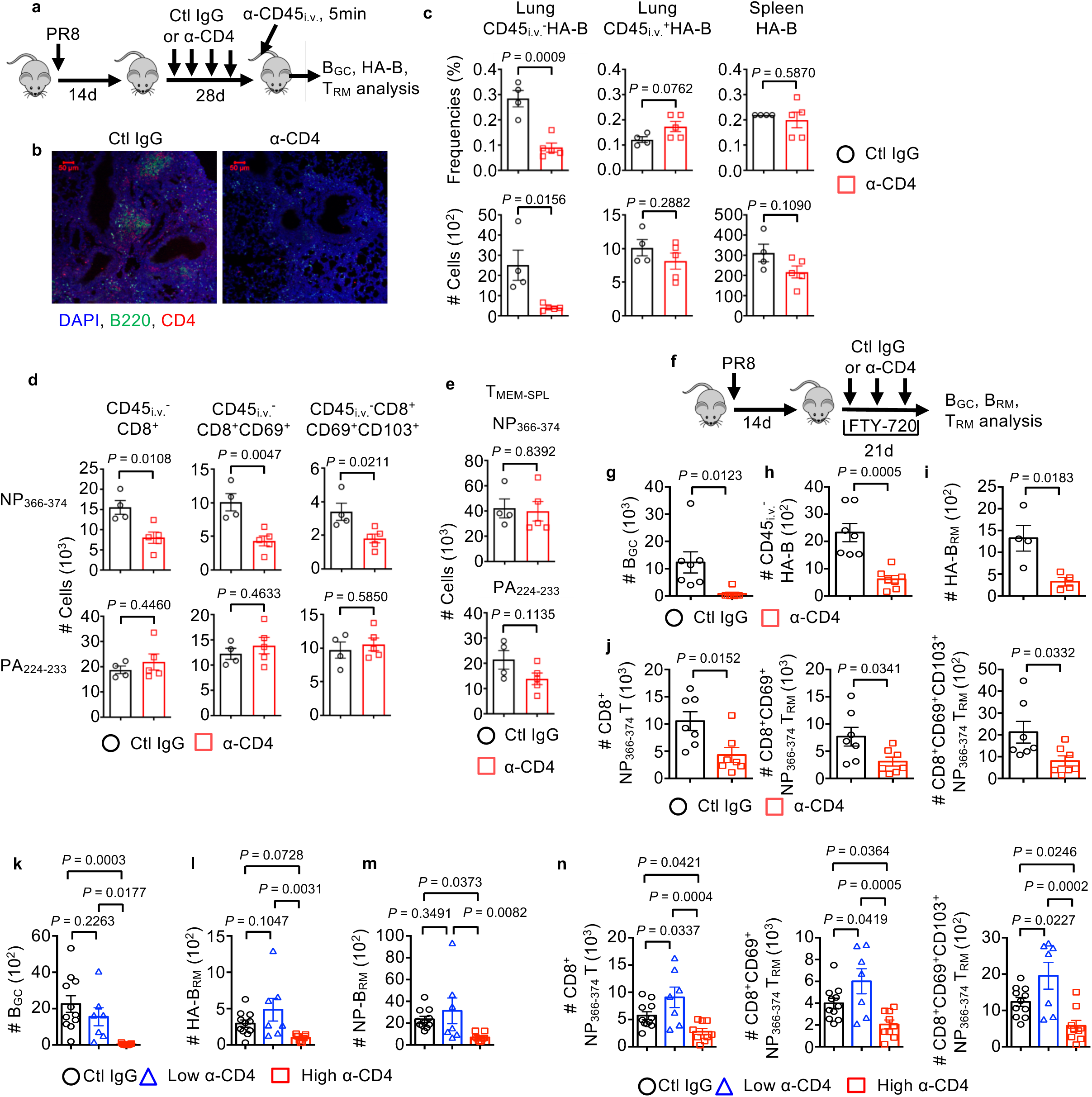
Lung CD4^+^ T cells deliver localized help for the development of tissue-resident B and CD8^+^ T cells. **a-e**, WT mice were infected with PR8 strain of influenza virus and treated with control (ctl) IgG or α-CD4 starting at 14 d.p.i. Mice were injected with α-CD45 intravenously (i.v.) 5 min before sacrifice at 42 d.p.i. **a**, Experimental scheme. **b**, Representative confocal images of iBALT in the lung. Lung sections were stained with α-CD4 (red), α-B220 (green) and DAPI (blue). **c**, Frequencies and cell number of influenza HA-specific B cells (HA-B) in the lung tissue (CD45_i.v._^-^B220^+^GL7^-^HA^+^), lung blood vessels (CD45_i.v._^+^B220^+^GL7^-^HA^+^) and spleen (B220^+^GL7^-^HA^+^). **d**, Lung tissue CD8^+^, CD8^+^CD69^+^ or CD8^+^CD69^+^CD103^+^ NP_366-374_ or PA_224-233_ T_RM_ cells were enumerated. **e**, Splenic CD8^+^ NP_366-374_ or PA_224-233_ memory cells (T_MEM-SPL_) were enumerated. **f-j**, WT mice were infected with PR8 and treated with ctrl IgG or α-CD4 (starting at 14 d.p.i.) in the presence of daily injection of FTY-720 (starting at 13 d.p.i.). **f**, Schematic of experimental design. **g**, B_GC_ cell numbers were enumerated by flow cytometry. **h, i**, Total HA-specific B cells (total HA-B) (**h**) or HA-specific tissue-resident memory B cells (HA-B_RM_: CD45_i.v._^-^B220^+^GL7^-^IgD^-^IgM^-^HA^+^CD38^+^) (**i**) were enumerated. **j**, Lung tissue CD8^+^, CD8^+^CD69^+^ or CD8^+^CD69^+^CD103^+^ NP_366-374_ or PA_224-233_ T_RM_ cells were enumerated. **k-n**, WT mice were infected with PR8 and received ctl IgG, high or low dose of α-CD4. Cell number of B_GC_ (**k**), HA-specific B_RM_ (**l**) and NP-specific B_RM_ cells (**m**) in the lung tissue. **n**, Lung tissue CD8^+^, CD8^+^CD69^+^ or CD8^+^CD69^+^CD103^+^ NP_366-374_ T_RM_ cells were enumerated. In **b-e** were the representative data from at least two independent experiments (4-5 mice per group). In **g-h and j-n**, data were pooled from two (**g, h** and **j**) or three (**k**-**n**) independent experiments (2-5 mice per group). *P* values were calculated by unpaired two-tailed Student’s t-test in c-j. *P* values in k-n were analyzed by one-way ANOVA.

We next examined influenza-specific CD8^+^ memory T cell responses in the lungs and spleens following CD4^+^ T cell depletion. To do so, we checked lung CD8^+^ memory T cell responses against two dominant influenza MHC-I H-2D^b^-restricted epitopes, nucleoprotein peptide 366-374 (NP_366-374_) and polymerase peptide 224-233 (PA_224-233_) through tetramer staining at 42 d.p.i. 40. It has been shown before that CD8^+^ memory T cells against NP_366-374_ or PA_224-233_ epitope exhibit distinct phenotypic and recall properties ^40, 48, 49, 50^. Specifically, lung CD8^+^ NP_366-374_ memory T cells are highly protective and dominate over the CD8^+^ PA_224-233_ memory T cells in the secondary recall expansion upon re-challenge with heterotypic influenza virus ^40, 48, 49, 50^. We found that late CD4^+^ T cell depletion did not affect lung circulating or tissue CD8^+^ PA_224-233_ memory T cell population (Fig. 1d and Extended Fig. 1f, g). However, late CD4^+^ T cell depletion caused significant decrease of the magnitude of CD8^+^ NP_366-374_ memory T cells in the lung tissue compartment but not in the circulation (Fig. 1d and Extended Fig. 1f, g). The magnitudes of parenchymal CD8^+^ CD69^+^ NP_366-374_ T_RM_ or CD8^+^ CD69^+^ CD103^+^ NP_366-374_ T_RM_ cells were also diminished following late CD4^+^ T cell depletion (Fig. 1d and Extended Fig. 1h). Notably, late CD4^+^ T cell depletion did not affect the percentages of CD103^+^ cells within the CD8^+^ CD69^+^ NP_366-374_ T_RM_ population (Extended Fig. 1i), which is in contrast with the results observed following CD4^+^ T cell depletion before influenza infection ^30^. Contrary to the diminished magnitude of lung CD8^+^ NP_366-374_ T_RM_ cells, late CD4^+^ T cell depletion did not decrease CD8^+^ NP_366-374_ or CD8^+^ PA_224-233_ memory T cells (T_MEM_) in the lung vasculature or spleen (Fig. 1e and Extended Fig. 1g). Thus, these data suggest that continuous CD4^+^ T cell help following viral clearance is required for the persistence of optimal B and CD8^+^ T cell responses (against a dominant protective epitope) in the lung at the memory stage.

It is possible that CD4^+^ T cells may provide the “help” either in the circulation or in the lungs for the generation of optimal tissue memory B and CD8^+^ T cells. To determine whether lung tissue CD4^+^ T cells can provide the “local help” for lung memory B and CD8^+^ T cell generation, we infected WT mice with PR8 virus and depleted CD4^+^ T cells at 14 d.p.i. in the presence of FTY720 (Fig. 1f), a chemical that blocks lymphocyte migration ^51^. We confirmed that FTY720 treatment drastically diminished T and B cell circulation in the blood (Extended Fig. 2a). The depletion of CD4^+^ T cells abolished B_GC_ cell development, total HA-specific B cells (HA-B) and strikingly HA-specific B_RM_ (HA-B_RM_) cells (identified as CD45_i.v._^-^ B220^+^ GL7^-^ IgD^-^ IgM^-^ CD38^+^ HA^+^, Extended Fig. 1j) in the lungs (Fig. 1 g-i) even in the presence of FTY720. CD4^+^ T cell depletion also diminished lung tissue CD8^+^ NP_366-374_-specific T_MEM_, CD8^+^ CD69^+^ T_RM_ and CD8^+^ CD69^+^ CD103^+^ T_RM_ cells (Fig. 1j). Similar data were observed following CD4^+^ T cell depletion in the presence of FTY720 during H3N2 X31 influenza virus infection (Extended Fig. 2 b-e). Thus, lung CD4^+^ T cells can provide *in situ* “help” to local B and CD8^+^ T cells for the optimal development of B_GC_, B_RM_ and CD8^+^ T_RM_ cells following viral clearance.

**Fig. 2.**
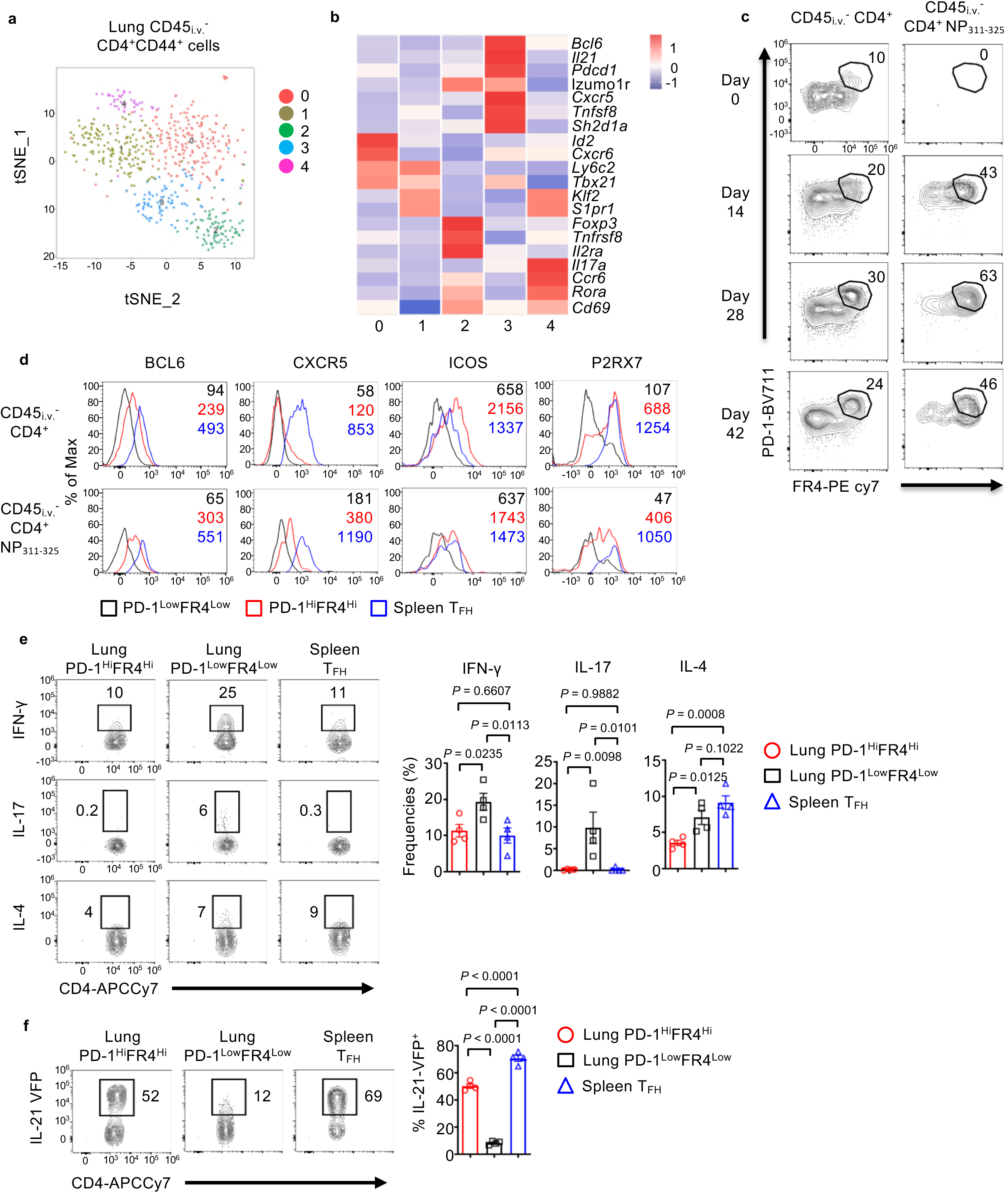
Identification of a population of T_FH_-like cells in the lung tissue. WT (**a-e**) or IL-21-VFP reporter (**f**) mice were infected with PR8. **a**, tSNE plot of scRNAseq analysis of sorted lung CD45_i.v._^-^CD4^+^CD44^Hi^ cells (pooled from 5 mice) at 28 d.p.i. **b**, Heat map of indicated genes in each cluster from scRNAseq data. **c**, Kinetics of the percentages of PD-1^Hi^FR4^Hi^ population in lung tissue total CD4^+^ or influenza NP-specific (NP_311-325_) CD4^+^ T cells. **d**, Expression of T_FH_ cell-associated markers in lung total or influenza-specific PD-1^Hi^FR4^Hi^, PD-1^Low^FR4^Low^ or splenic T_FH_ (CD4^+^CD44^Hi^PD-1^+^CXCR5^+^) cells at 28 d.p.i. **e**, Expression of IFN-γ, IL-17 or IL-4 by lung PD-1^Hi^FR4^Hi^, PD-1^Low^FR4^Low^ or spleen T_FH_ cells were identified by intracellular staining at 28 d.p.i**. f**, IL-21-VFP expression in lung CD4^+^ PD-1^Hi^FR4^Hi^, PD-1^Low^FR4^Low^ or spleen T_FH_ at 28 d.p.i. In **c-f**, representative plots, histograms and graphs were from at least two independent experiments (4 mice per group). *P* values in e and f were analyzed by one-way ANOVA.

We next sought to examine whether lung tissue CD4^+^ T cells are sufficient for the generation of B_GC_, B_RM_ and CD8^+^ T_RM_ responses following the clearance of the primary infection, in the absence of circulating CD4^+^ T cells. To do so, we injected low or high doses of α-CD4 into PR8-infected WT mice at 14 d.p.i. Low dose α-CD4 treatment largely ablated CD4^+^ T cells in the circulating blood, but not in the lung parenchyma, while high dose α-CD4 injection depleted both circulating and lung parenchymal CD4^+^ T cells (Extended Fig. 2 f-h). Strikingly, the high dose, but not the low dose, of α-CD4 treatment diminished the magnitude of lung B_GC_, influenza NP (nucleoprotein)-specific B_RM_ and CD8^+^ NP_366-374_ T_RM_ (including CD69^+^ T_RM_ or CD69^+^CD103^+^ T_RM_) cells (Fig. 1 k-n). Taken together, our data suggest that lung tissue CD4^+^ T cells, rather than CD4^+^ T cells in the circulation, provide late “local help” for the optimal generation of B_GC_, B_RM_ and CD8^+^ T_RM_ responses following influenza infection.

### Identification of a population of T_FH_-like cells in the lung

We next sought to identify the lung CD4^+^ T cell populations that may mediate the local help to B and/or CD8^+^ T cells. To do so, we performed single cell (sc) RNA-seq on CD45_i.v._^-^ lung parenchymal CD4^+^ T cells at 28 d.p.i. Hierarchical clustering analysis identified 5 separated CD4^+^ T cell populations within the lung parenchyma (Fig. 2a and Extended Fig. 3 a, b). These cell populations included Th1 effector/memory-like cells (expressing high levels of *Cxcr6/Tbx21* (T-bet), cluster 0), T cells recently entering the lungs or circulating effector memory T cells (expressing high *Klf2/S1pr1*, cluster 1), regulatory T cells (Treg, expressing *Foxp3*, cluster 2), Th17-like (expressing *Il17/Ccr6/Rora*, cluster 4) and a cluster of T cells exhibiting features of T_FH_ cells (expressing *Bcl6/Il21*, cluster 3) (Fig. 2b and Extended Fig. 3 a-c). Cluster 3 CD4^+^ T cells also expressed higher levels of T_FH_–associated surface molecules *Pdcd1*(PD-1), *Cxcr5* and *Izumo1r* (FR4) (Fig. 2b).

**Fig. 3.**
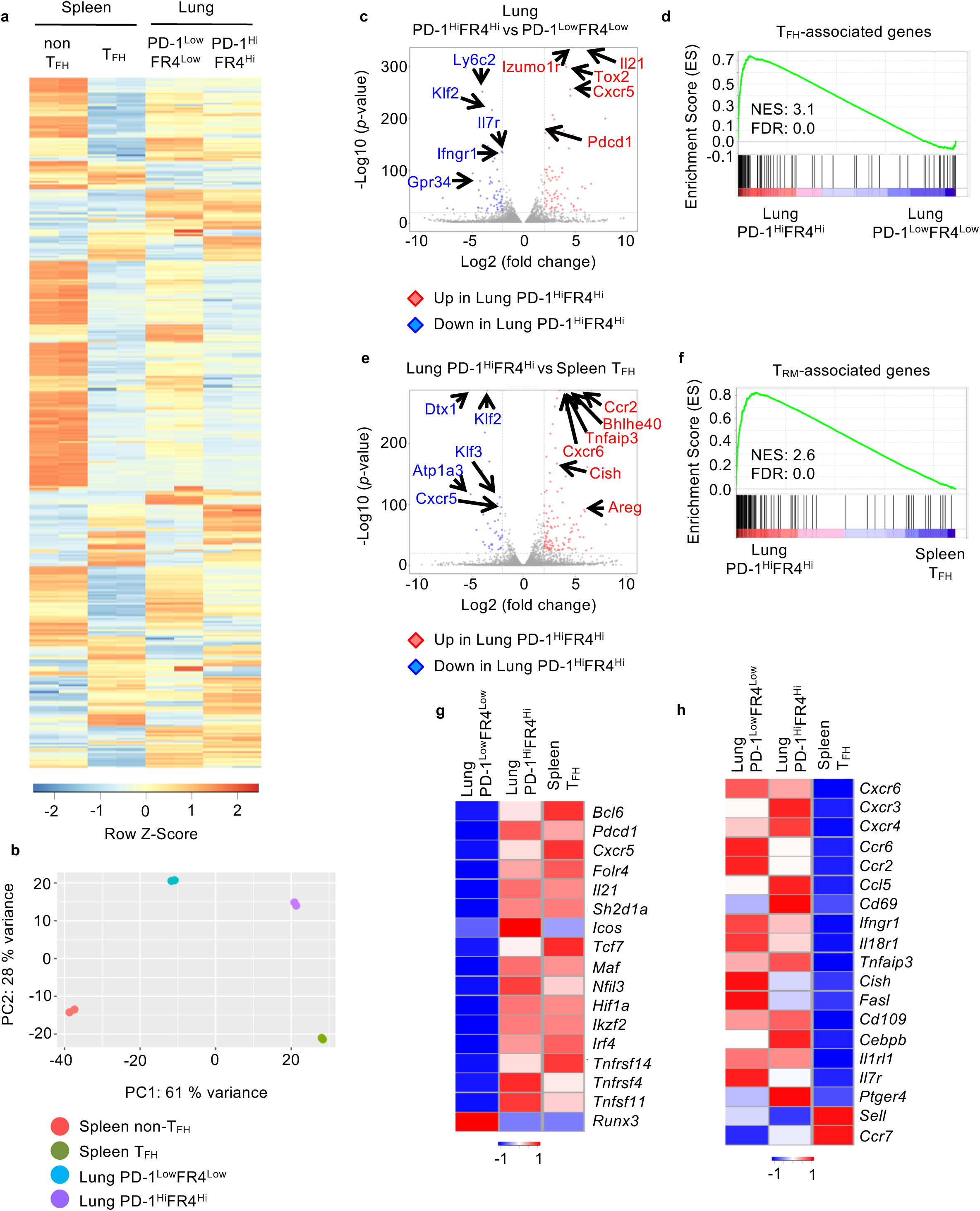
Transcriptional profiling reveals PD-1^Hi^FR4^Hi^ cells exhibit both T_FH_ and T_RM_ gene signatures. **a-f**, WT mice were infected with PR8. Lung PD-1^Hi^FR4^Hi^ or PD-1^Low^FR4^Low^ CD4^+^ T cells and splenic T_FH_ or non-T_FH_ cells were sorted following exclusion of GITR^Hi^ Treg cells and RNA-seq analysis was performed at 28 d.p.i. **a**, Heatmap expression of differentially expressed genes among lung PD-1^Hi^FR4^Hi^, PD-1^Low^FR4^Low^ CD4^+^ T cells and splenic T_FH_ or non-T_FH_ cells. **b**, Principle component analysis of RNA-seq data of lung PD-1^Hi^FR4^Hi^, PD-1^Low^FR4^Low^ CD4^+^ T cells and splenic T_FH_ or non-T_FH_ cells. **c**, Volcano plot of RNA-seq analysis of lung PD-1^Hi^FR4^Hi^ or PD-1^Low^FR4^Low^ CD4^+^ T cells. **d**, GSEA of the core T_FH_ signature genes in lung CD4^+^ PD-1^Hi^FR4^Hi^ and CD4^+^ PD-1^Low^FR4^Low^ cells. **e**, Volcano plot of RNA-seq analysis on lung PD-1^Hi^ FR4^Hi^ CD4^+^ T and splenic T_FH_ cells. **f**, GSEA of the core tissue-residency signature genes of T_RM_ cells in lung PD-1^Hi^FR4^Hi^ and splenic T_FH_ cells. **g-h**, WT mice were infected with PR8. Lung PD-1^Hi^FR4^Hi^ or PD-1^Low^FR4^Low^ CD4^+^ T cells and splenic T_FH_ cells were sorted at 28 d.p.i. Nanostring analysis on 560 immune-associated genes was performed. The expression of T_FH_-associated genes (**g**) or tissue-residency associated genes (**h**) in the three cell populations was depicted. For RNA-seq, data were from duplicates of pooled samples (n = 15). For nanostring analysis, data were from pooled samples (n = 10).

Flow cytometry analysis identified a population of total CD4^+^ or influenza-specific CD4^+^ NP_311-_ 325 PD-1^Hi^FR4^Hi^ T cells that developed following influenza infection and expressed T_FH_-associated markers including BCL6, ICOS and P2RX7, but relatively lower levels of CXCR5 compared to splenic T_FH_ cells (Fig. 2 c, d). Similar to splenic T_FH_ cells, lung CD4^+^ PD-1^Hi^FR4^Hi^ T cells expressed low levels of IFN-γ and IL-17 (Fig. 2e). However, splenic T_FH_ cells expressed significantly higher IL-4 than lung CD4^+^ PD-1^Hi^FR4^Hi^ T cells (Fig. 2 e). We next examined whether lung CD4^+^ PD-1^Hi^FR4^Hi^ T cells expressed the T_FH_ signature cytokine IL-21 following influenza virus infection using the IL-21-VFP reporter mice ^52^. Lung CD4^+^ PD-1^Low^ FR4^Low^ T cell population expressed modest levels of IL-21, but the lung CD4^+^ PD-1^Hi^FR4^Hi^ T cells expressed high levels of IL-21, almost comparable to those of splenic T_FH_ cells (Fig. 2f). These data suggest that there is a lung tissue CD4^+^ T cell population, transcriptionally and phenotypically resembling T_FH_ cells in the secondary lymphoid organs.

### Transcriptional profiling reveals lung T_FH_-like cells exhibit T_RM_ gene signature

To gain more insights into the phenotype and identity of lung T_FH_-like cells, we sought to compare the transcriptional signatures of lung T_FH_-like cells to splenic T_FH_, non-T_FH_ and lung non-T_FH_-like cell signatures. Foxp3^+^ Treg cells expressed FR4, GITR and PD-1 (albeit their PD-1 levels were lower than T_FH_-like cells) (Fig. 2b and Extended Fig. 3c and 4a). To minimize the potential contribution of Treg cells on the transcriptional profiles of lung T_FH_-like cells, we excluded splenic or lung GITR^+^ CD4^+^ T cells, which were mostly Foxp3^+^ cells, in our sorting (Extended Fig. 4a). We then sorted splenic non-T_FH_ (CD4^+^CD44^Hi^GITR^-^CXCR5^Low^PD-1^Low^), splenic T_FH_ (CD4^+^CD44^Hi^GITR^-^CXCR5^Hi^PD-1^Hi^), lung non-T_FH_-like cells (CD45_i.v._^-^ CD4^+^CD44^Hi^GITR^-^PD-1^Low^FR4^Low^) and lung T_FH_-like cells (CD45_i.v._^-^ CD4^+^CD44^Hi^GITR^-^ PD-1^Hi^ FR4^Hi^) cells and performed RNA-seq analysis.

**Fig. 4.**
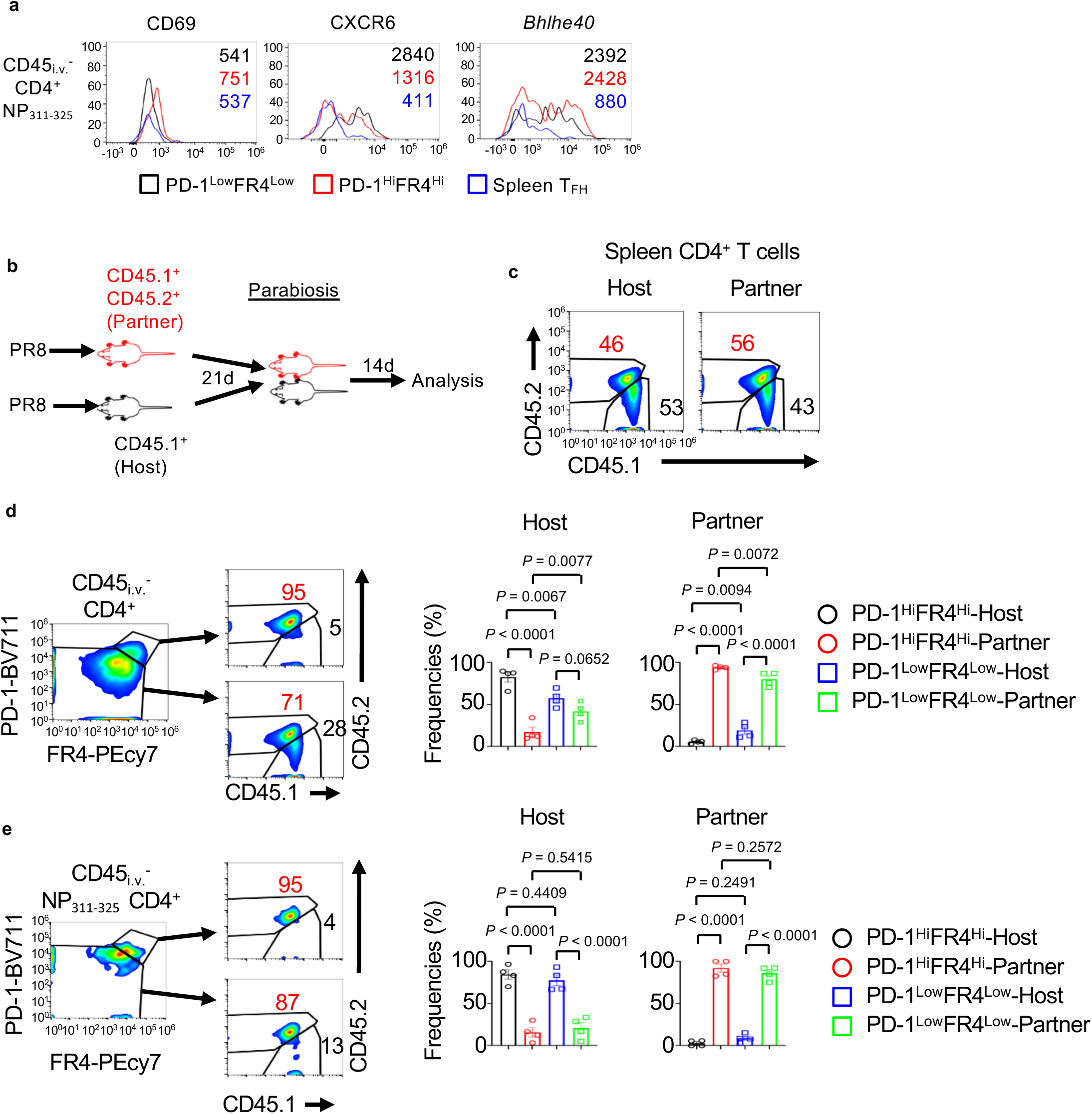
Lung PD-1^Hi^FR4^Hi^ cells are tissue resident. **a**, WT mice were infected with PR8. The expression of CD69, CXCR6 and *Bhlhe40* in lung CD4^+^ PD-1^Hi^FR4^Hi^ or CD4^+^ PD-1^Low^FR4^Low^ NP_311-325_ T cells or splenic T_FH_ cells at 28 d.p.i. **b-e**, CD45.1^+^ (Host) or CD45.1^+^ CD45.2^+^ (Partner) WT mice were infected with PR8. Parabiosis surgery was performed at 21 d.p.i. Mice were sacrificed 14 days later for analysis. **b**, Schematics of parabiosis experiments. **c**, Composition of Host-derived or Partner-derived CD4^+^ T cells in the spleens of the parabionts. **d**, Frequencies of Host-derived or Partner-derived cells in the lung PD-1^Hi^FR4^Hi^ or PD-1^Low^FR4^Low^ total CD4^+^ T cell compartment. **e**, Frequencies of Host-derived or Partner-derived cells in influenza-specific lung CD4^+^ PD-1^Hi^FR4^Hi^ or CD4^+^ PD-1^Low^FR4^Low^ NP_311-325_ T cell compartment. In **a**, the representative histograms were from at least two independent experiments (3-4 mice per group). Parabiosis data were pooled from two different experiments (two pairs per experiment). *P* values in d and e were analyzed by one-way ANOVA.

Differential gene expression and principal component analysis revealed that those four different CD4^+^ T cell populations have distinct gene expression patterns, although lung CD4^+^ PD-1^Hi^FR4^Hi^ cells were more distinct from splenic non-T_FH_ cells relative to splenic T_FH_ or lung CD4^+^ PD-1^Low^FR4^Low^ cells (Fig. 3a, b and Extended Fig. 4b). When directly compared to lung CD4^+^ PD-1^Low^FR4^Low^ cells, lung CD4^+^ PD-1^Hi^FR4^Hi^ cells highly expressed T_FH_-associated genes including *Il21, Tox2* and *Pdcd1* (Fig. 3c). Indeed, Gene Set Enrichment Analysis (GSEA) showed that lung CD4^+^ PD-1^Hi^FR4^Hi^ cells had enrichment of T_FH_-associated genes ^53^ relative to CD4^+^ PD-1^Low^FR4^Low^ cells (Fig. 3d). Conversely, lung CD4^+^ PD-1^Low^FR4^Low^ cells expressed higher levels of *Ly6c* and *Il7r*, and showed enhanced enrichment of genes in TGF-β, hypoxia and Notch signaling relative to PD-1^Hi^FR4^Hi^ cells (Fig. 3c and Extended Fig. 4c). When compared to splenic T_FH_ cells, lung CD4^+^ PD-1^Hi^FR4^Hi^ cells showed increased expression of genes associated with tissue migration, retention and function including *Ccr2, Bhlhe40* and *Cxcr6* (Fig. 3e) ^54, 55^. GSEA analysis revealed that lung CD4^+^ PD-1^Hi^FR4^Hi^ had significant enrichment of T_RM_-associated genes relative to splenic T_FH_ cells (Fig. 3f) ^56^. Lung CD4^+^ PD-1^Hi^FR4^Hi^cells also had higher expression of genes associated with IL-2/STAT5, NF-κB and interferon signaling, whereas splenic T_FH_ cells had higher enrichment of Myc and PI3K-mTOR signaling (Extended Fig. 4d). Thus, these RNA-seq analyses indicate that lung CD4^+^ PD-1^Hi^FR4^Hi^ cells exhibit transcriptional signatures of both T_FH_ cells and T_RM_ cells.

To confirm these observations, we sorted splenic T_FH_, lung CD4^+^ PD-1^Hi^FR4^Hi^ or lung CD4^+^ PD-1^Low^FR4^Low^ cells and performed Nanostring analysis of 560-immune associated genes without the need of RNA amplification ^40^. Compared to lung CD4^+^ PD-1^Low^FR4^Low^ cells, splenic T_FH_ and lung CD4^+^ PD-1^Hi^FR4^Hi^ cells expressed higher levels of T_FH_-associated genes including *Bcl6, Sh2d1a* and *Tcf7* (Fig. 3g). Compared to splenic T_FH_ cells, both PD-1^Hi^ FR4^Hi^ and PD-1^Low^ FR4^Low^ lung CD4^+^ T cell populations had enhanced expression of genes associated with tissue migration and residency, but diminished expression of lymphoid migration or retention molecules *Sell* (CD62L) and *Ccr7* (Fig. 3h). Altogether, these data suggest that lung CD4^+^ PD-1^Hi^FR4^Hi^ cells exhibit a “hybrid” gene signature of both conventional T_FH_ cells and T_RM_ cells.

### Lung CD4^+^ PD-1^Hi^ FR4^Hi^ T cells are tissue-resident

Given that lung CD4^+^ PD-1^Hi^FR4^Hi^ cells showed gene signatures of tissue residency, we sought to determine whether these cells are indeed tissue-resident. Using flow cytometry, we confirmed that PD-1^Hi^FR4^Hi^ cells expressed higher levels of CD69, CXCR6 and *Bhlhe40*, molecules associated with T cell migration, retention and function in the respiratory mucosal tissue, compared to splenic T_FH_ cells (Fig. 4a). We then performed parabiosis experiments and joined the circulation of PR8-infected CD45.1^+^ and CD45.1^+^ CD45.2^+^ congenic mice at 21 d.p.i. (Fig. 4b). We examined CD4^+^ T cell exchange between the two parabionts after 2 weeks of parabiosis. Close to 40-60 % of splenic CD4^+^ T cells in parabiont hosts were derived from their parabiont pair (Fig. 4c), suggesting the successful exchange of circulating CD4^+^ T cells between the parabionts. Within lung i.v. CD45 antibody (Ab) protected tissue CD4^+^ T cell compartment, PD-1^Hi^FR4^Hi^ total CD4^+^ or antigen (Ag)-specific CD4^+^ NP_311-325_ T cells exhibited limited exchange between the two parabionts (Fig. 4d, e), suggesting that these cells are mostly tissue-resident. Of note, most of Ag-specific CD4^+^ NP_311-325_ T cells are tissue-resident (Fig. 4e), while total lung CD4^+^ PD-1^Low^FR4^Low^ T cells showed higher circulating rate than those of CD4^+^ PD-1^Hi^FR4^Hi^ cells (Fig. 4 d). Thus, lung CD4^+^ PD-1^Hi^FR4^Hi^ cells are lung tissue-resident T cells exhibiting both T_FH_ and T_RM_ features. Based on their gene signature, protein expression, cytokine production and tissue residency property, we termed these cells as tissue-resident T helper (T_RH_) cells.

### T_RH_ responses require B cells and lung tertiary lymphoid structure formation

T_FH_ generation requires the presence of B cells ^57^. We therefore examined whether lung B cells are required for the generation of T_RH_ cells following influenza infection. To do so, we infected WT mice with PR8 and then depleted B cells with α-CD20 treatment in the presence of FTY-720 to block T/B cell migration (Fig. 5a). We validated that α-CD20 treatment depleted lung B cells (Extended Fig. 5a). We then determined splenic T_FH_ and lung T_RH_ responses with or without α-CD20 treatment at 28 d.p.i. Consistent with previous findings ^58^, B cell depletion following α-CD20 treatment diminished splenic T_FH_ responses (Fig. 5b). α-CD20 treatment also impaired lung T_RH_ responses, but not lung non-T_RH_ (PD-1^Low^FR4^Low^) responses. Thus, lung B cells are required for the development of T_RH_ responses. Tertiary lymphoid structures (iBALT) form in the lung that consist of aggregated B, T and dendritic cells (DCs) following influenza viral clearance ^37, 46, 47^. IL-21 expressing lung CD4^+^ T cells were found in the iBALT from PR8-infected IL-21-VFP reporter mice (Extended Fig. 5b), suggesting that iBALT formation may be required for optimal T_RH_ responses following influenza virus clearance. To this end, we infected WT mice with PR8 and then injected the mice with lymphotoxin beta receptor Ig fusion protein (LTβR-Ig) in the presence of FTY-720 to deplete iBALTs in the lungs ^59, 60^ (Fig. 5a). We confirmed that LTβR-Ig injection diminished lung iBALT formation and the magnitude of lung parenchyma B cell expansion (Fig. 5c and Extended Fig. 5c). LTβR-Ig injection did not affect splenic T_FH_ responses, but significantly impaired influenza-specific lung T_RH_ but not non-T_RH_ responses (Fig. 5d). Thus, these data indicate that lung tissue B cells and iBALT formation is required for the optimal generation of lung T_RH_ responses.

**Fig. 5.**
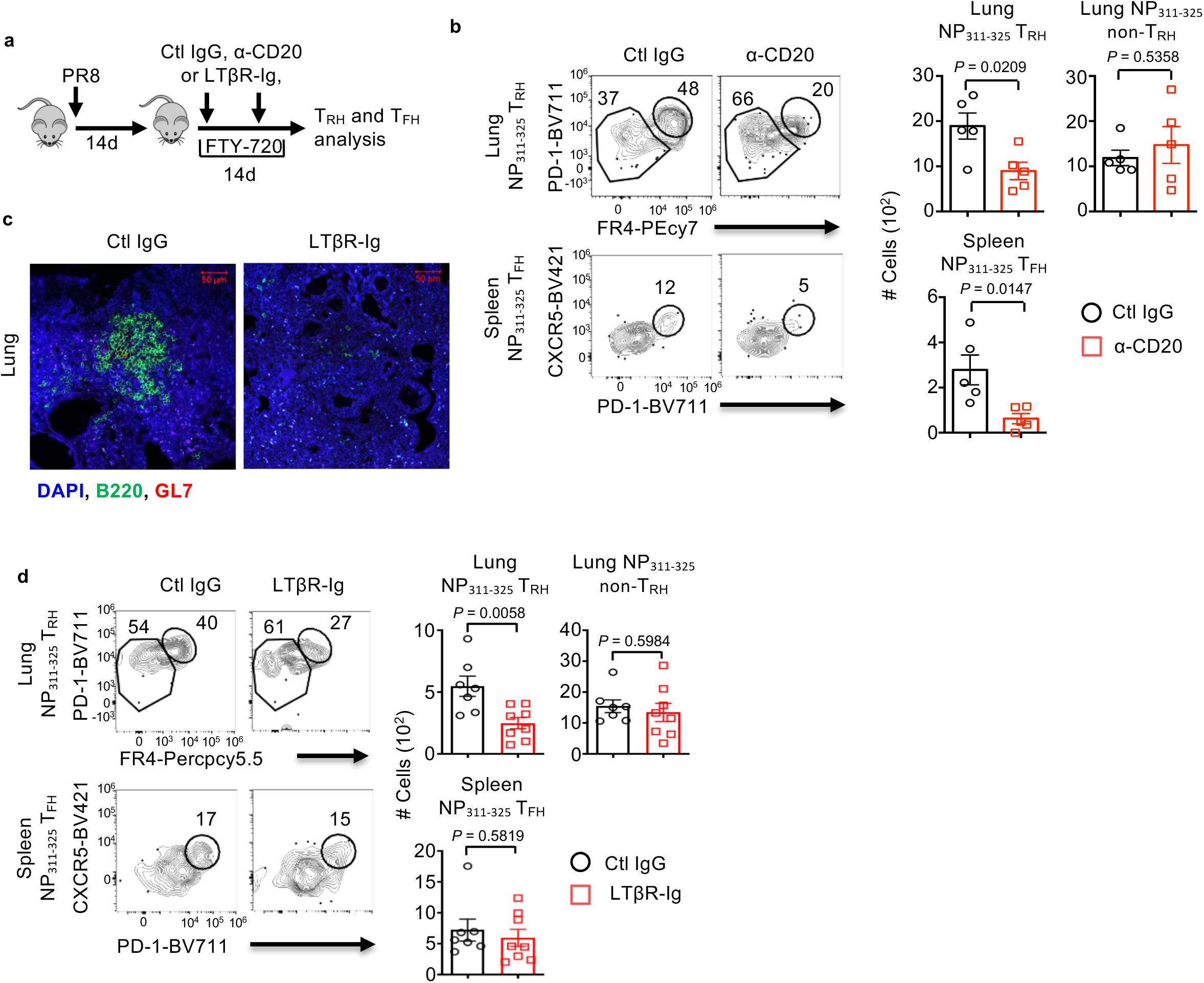
Optimal T_RH_ formation requires lung B cells and iBALT formation. WT (**a-d**) mice were infected with PR8 and treated with ctl IgG, α-CD20 (**b**) or LTβR-Ig (**c** and **d**) (weekly starting at 14 d.p.i.) in the presence of daily injection of FTY-720 (13-27 d.p.i.). **a**, Experimental scheme. **b** and **d**, Representative dot plot and cell numbers of influenza-specific NP_311-325_ lung T_RH,_ lung non-T_RH_ or splenic T_FH_ cells. **c**, Representative image from lung tissue section stained with B220/GL-7 following ctl IgG or LTβR-Ig treatment. In **c**, the representative image was from at least two independent experiments (3-4 mice per group). In **b**, representative data were from at least two independent experiments (4-5 mice per group). In **d**, data were pooled from two independent experiments (3-4 mice per group). *P* values were calculated by unpaired two-tailed Student’s t-test.

### Optimal T_RH_ responses depend on both T_FH_ and T_RM_ transcription factors

We next sought to investigate the molecular mechanisms regulating lung T_RH_ cell development following influenza virus infection. We first examined whether lung T_RH_ cell development is dependent on the master transcription factor of T_FH_ cells, BCL6 ^61, 62^. To do so, we infected WT (*Bcl6*^*fl/fl*^) or *Bcl6*^*ΔT*^ mice with PR8 virus and examined total and influenza-specific T_RH_ or non-T_RH_ cells in the lung tissue at 28 d.p.i. We found that T cell-specific BCL6 deficiency greatly diminished both the frequencies and the magnitude of lung T_RH_ responses, but not those of non-T_RH_ responses (Fig. 6a-c). Consistent with the literatures ^57, 63^, T cell-specific BCL6 deficiency also diminished splenic T_FH_ responses (Fig. 6d, e).

**Fig. 6.**
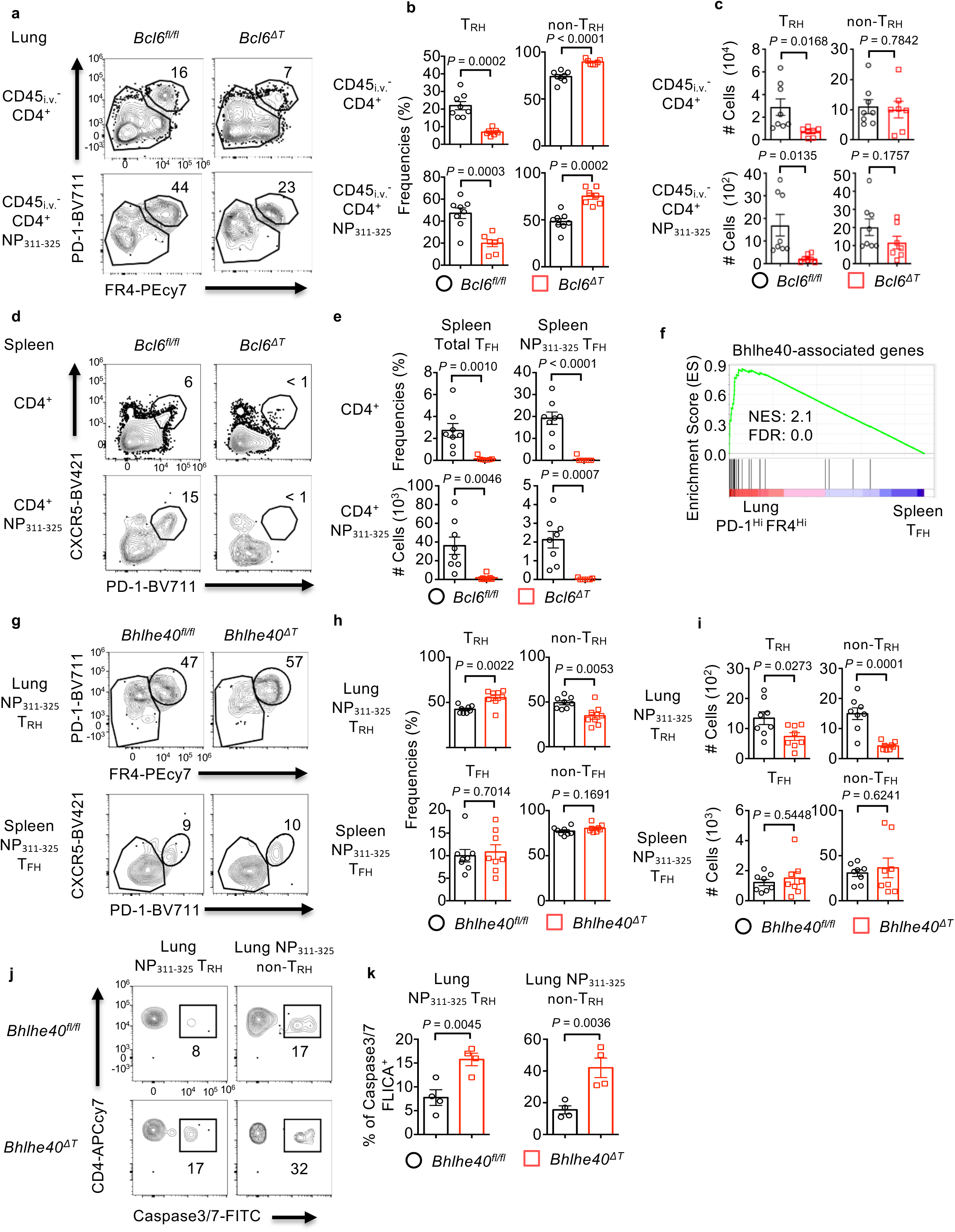
Both BCL6 and Bhlhe40 are required for optimal lung T_RH_ responses. **a-e**, *Bcl6*^*fl/fl*^ or *Bcl6*^*ΔT*^ mice were infected with PR8. Representative dot plot (**a**), percentages (**b**) and cell numbers (**c**) of T_RH_ or non-T_RH_ in lung CD45_i.v._^-^ total CD4^+^ or CD45_i.v._^-^ influenza-specific CD4^+^ NP_311-325_ T cells at 28 d.p.i. **d-e**, Representative dot plot (**d**) and percentage (top row) or cell numbers (bottom row) (**e**) of splenic total T_FH_ or NP_311-325_ T_FH_ at 28 d.p.i. **f**, GSEA of the *Bhlhe40*-associated genes in lung T_RH_ (PD-1^Hi^FR4^Hi^) and spleen T_FH_ cells. **g-k**, *Bhlhe40*^*fl/fl*^ or *Bhlhe40*^*ΔT*^ mice were infected with PR8. **g-i**, Representative dot plot (**g**), percentages (**h**) or cell numbers (**i**) of influenza-specific NP_311-325_ lung T_RH_, lung non-T_RH_, splenic T_FH_ or splenic non-T_FH_ cells at 28 d.p.i. **j-k** Representative dot plot (**j**) or percentages (**k**) of active caspase 3/7-FLICA^+^ cells in lung NP_311-325_ T_RH_ or non-T_RH_ cells at 28 d.p.i. In **a-e** and **g-i**, data were pooled from two independent experiments (3-4 mice per group). In **j-k**, representative data were from at least two independent experiments (4 mice per group). *P* values were calculated by unpaired two-tailed Student’s t-test.

We have demonstrated before that Bhlhe40 is critical for the development of tissue-resident CD8^+^ T cell responses ^55^. Since lung T_RH_ and non-T_RH_ cells expressed high levels of *Bhlhe40* relative to splenic T_FH_ cells (Fig 4a), we investigated the roles of Bhlhe40 in lung T_RH_ responses relative to splenic T_FH_ cells. Consistent with high Bhlhe40 expression in T_RH_ cells, lung T_RH_ cells were enriched with Bhlhe40-associated genes ^55^ compared to splenic T_FH_ cells (Fig. 6f). We then infected WT (*Bhlhe40*^*fl/fl*^) or *Bhlhe40*^*ΔT*^ mice with PR8 and examined total and influenza-specific T_RH_ or non-T_RH_ cells at 28 d.p.i. T cell-specific Bhlhe40 deficiency modestly increased the frequencies of T_RH_ cells relative to non-T_RH_ cells within the influenza-specific NP_311-325_ CD4^+^ T cell population, but not in the total lung CD4^+^ T cell population (Fig. 6g, h and Extended Fig. 5 e). Bhlhe40-deficiency in T cells significantly diminished total and influenza-specific CD4^+^ T cells in the lung tissue, but not in the spleen (Extended Fig. 5 f, g). These data suggest that Bhlhe40 is required for the establishment of the overall lung-resident CD4^+^ T cell population following influenza infection, as was observed with the lung-resident CD8^+^ T cells ^55^. As the result, the magnitude of both T_RH_ and non-T_RH_ cells in the lungs were significantly decreased (Fig. 6i). In contrast, Bhlhe40 deficiency did not alter either the frequencies or the magnitude of the splenic T_FH_ response (Fig. 6g-i and Extended 5 d, e). Previously, we have reported that Bhlhe40 is required for the survival CD8^+^ T cells in non-lymphoid tissues ^55^. Consistent with that observation, Bhlhe40 deficiency resulted in enhanced cellular apoptosis in both lung T_RH_ and non-T_RH_ cells (Fig. 6j, k). These data indicate that Bhlhe40 is likely not important for the acquisition of “T_FH_-like” features in T_RH_ cells, but is vital in sustaining T_RH_ cell survival in the respiratory mucosal tissues. Taken together, the optimal formation of lung T_RH_ cells requires transcription factors involved in both T_FH_ (BCL6) and T_RM_ (Bhlhe40) development. Conversely, the formation of splenic T_FH_ cells is dependent on BCL6 but not Bhlhe40, and the development of lung PD-1^Low^FR4^Low^ cells (probably consist of conventional T_RM_ cells) is dependent on Bhlhe40 but not BCL6.

### T_RH_ cells assist the formation of protective B_GC_, B_RM_ and CD8^+^ T_RM_ responses

We hypothesize that T_RH_ cells are those cells mediating the effects of CD4^+^ T cell help on lung local B and CD8^+^ T cells. Consistent with that hypothesis, T cell-specific BCL6 deficiency leaded to diminished iBALT formation, B_GC_, B_RM_ and CD8^+^ NP_366-374_ T_RM_ responses (Extended Fig. 6 a-d). Furthermore, T cell-specific Bhlhe40 deficiency also resulted in diminished lung tissue B cells, B_RM_ and CD8^+^ T_RM_ responses, although it is possible that Bhlhe40 deficiency in CD8^+^ T cells directly contribute to the diminished CD8^+^ T_RM_ phenotype in these mice as shown before ^55^ (Extended Fig. 6 e-g).

To specifically determine whether T_RH_ cells are required for the development of memory B and CD8^+^ T cells in the lungs, we generated CD4 T cell -specific inducible BCL6-deficient mice (*Bcl6*^*ΔCD4 ERT2*^). We first confirmed that tamoxifen treatment efficiently caused gene recombination in CD4^+^ T cells, but only minimally in other lymphocytes in *Bcl6*^*ΔCD4 ERT2*^ mice (Extended Fig. 7a-c). We then infected WT (*Bcl6*^*fl/fl*^) or *Bcl6*^*ΔCD4 ERT2*^ mice with PR8 and inoculated tamoxifen daily from 12 to 16 d.p.i. (5X) to specifically ablate BCL6 in CD4^+^ T cells following influenza infection (Fig. 7a). To exclude the contribution of lymphoid organ T_FH_ cells in providing the “late” help to lung B and CD8^+^ T cells, we treated the mice with FTY720 daily to block T and B cell migration starting at 11 d.p.i. At the lung, the magnitude of CD45_i.v._^-^ CD4^+^, CD8^+^ T and B cells was decreased in inducible BCL6 ablated group (Extended Fig. 7d). As with the constitutive BCL6 deficiency in T cells, inducible CD4^+^ T cell-specific BCL6 ablation resulted in diminished T_RH_ but not non-T_RH_ responses in the lungs (Extended Fig. 7e). Strikingly, the ablation of lung T_RH_ responses significantly diminished B_GC_, influenza HA-specific B_RM_(HA-B_RM_) and NP-specific B_RM_ (NP-B_RM_) responses in the lungs (Fig. 7b-d). CD8^+^ NP_366-374_ T_RM_ responses were also significantly impaired following T_RH_ ablation (Fig. 7e). Together these data suggest that lung T_RH_ cells are important in assisting the development of local B and CD8^+^ memory responses *in situ* in the respiratory tract.

**Fig. 7.**
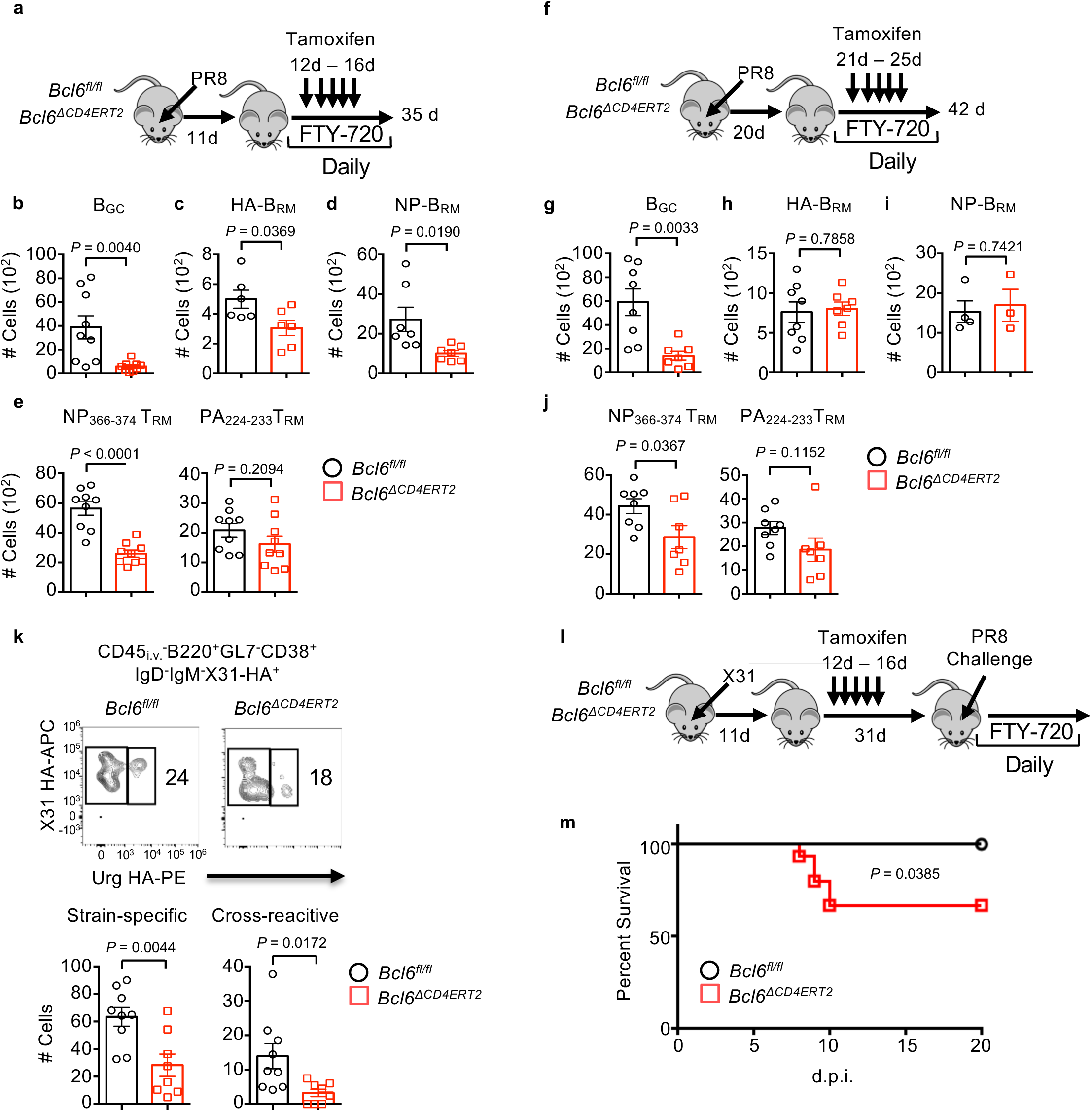
T_RH_ cells are required for the development of lung protective CD8^+^ T_RM_ and B cell immunity. **a-j**, *Bcl6*^*fl/fl*^ or *Bcl6*^*ΔCD4ERT2*^ mice were infected with PR8. **a-e**, Tamoxifen was administrated daily from 12-16 d.p.i. in the presence of daily FTY720 administration (11-34 d.p.i.). **a**, Schematics of experimental design. Cell numbers of B_GC_ (**b**), HA-specific B_RM_ (**c**), NP-specific B_RM_ (**d**), **e**, CD8^+^CD69^+^ NP_366-374_ T_RM_ or CD8^+^CD69^+^ PA_224-233_ T_RM_ at 35 d.p.i. **f-j**, Tamoxifen was administrated daily from 21-25 d.p.i. in the presence of daily FTY720 administration (20-41 d.p.i.). **f**, Schematics of experimental design. Cell numbers of B_GC_ (**g**), HA-specific B_RM_ (**h**), NP-specific B_RM_ (**i**), **j**, CD8^+^CD69^+^ NP_366-374_ T_RM_ or CD8^+^CD69^+^ PA_224-233_ T_RM_ at 42 d.p.i. **k**, *Bcl6*^*fl/fl*^ or *Bcl6*^*ΔCD4ERT2*^ mice were infected with X31 strain (H3N2) of influenza. Tamoxifen was administrated daily from 12-16 d.p.i. in the presence of daily FTY720 administration (11-34 d.p.i.). Representative dot plot (top) and cell numbers (bottom) of X31 strain-specific B_RM_ or cross-reactive HA-specific B_RM_ (to H3N2 A/Uruguay/716/07 strain) at 35 d.p.i. **l-m**, *Bcl6*^*fl/fl*^ (n = 11) or *Bcl6*^*ΔCD4ERT2*^ (n = 15) mice were infected with X31 and administered with tamoxifen from 12 to 16 d.p.i. Mice were re-challenged with PR8 at 42 d.p.i. in the presence of FTY720 (starting from 41d). **l**, Schematics of experimental design. **m**, Host mortality following PR8 challenge was monitored. In **a-h and j-k**, all data were pooled from two (**c, d, k, g, h** and **j**) or three (**b** and **e**) independent experiments (2-5 mice per group). In a-k, *P* values were calculated by unpaired two-tailed Student’s t-test. *P* value of survival study in m was calculated by Logrank test.

To examine the roles of T_RH_ cells in the maintenance of lung tissue B and CD8^+^ T cell responses, we treated WT or *Bcl6*^*ΔCD4 ERT2*^ mice with tamoxifen starting from 21 to 25 d.p.i. to ablate T_RH_ cell responses at the memory stage (Fig. 7f). We then examined B_GC_, B_RM_ and CD8^+^ T_RM_ responses at 42 d.p.i. (Fig. 7g-j). T_RH_ ablation at the memory stage did not lead to significant decrease of B_RM_ magnitude, but significantly impaired B_GC_ responses at 6 weeks post infection (Fig. 7g-i). These data suggest T_RH_ cells help to program lung B_RM_ cell development, but may not be directly required for their maintenance at the memory stage. However, T_RH_ cells are continuously needed for sustaining lung B_GC_ responses (Fig. 7g). Notably, these data are consistent with the data showing that B_GC_ responses are important for B_RM_ cell development in the first three weeks, but may not significantly contribute to lung B_RM_ cells after 20 d.p.i. ^5^. T_RH_ ablation at the memory stage also diminished CD8^+^ NP_366-374_ T_RM_ responses, suggesting that lung T_RH_ cells continuously provide “local help” for the maintenance of CD8^+^ NP_366-374_ T_RM_ cells (Fig. 7j). Taken together, our data suggest that T_RH_ cells are vital for programming lung B_RM_ cell development before 20 d.p.i., but are necessary for both the optimal development and maintenance of B_GC_ and CD8^+^ NP_366-374_ T_RM_ responses at the memory stage.

Previously, it was shown that lung memory B cells generated from local B_GC_ cells harbored high portions of cross-reactive B cells following influenza X31 virus infection (H3N2 virus) ^14^. To examine whether T_RH_ cells help to generate those cross-reactive B_RM_ cells, we infected WT or *Bcl6*^*ΔCD4ERT2*^ mice with X31 virus and injected the mice with tamoxifen in the presence of FTY720 as in Extended Fig. 7a. We then checked X31 strain-specific HA-B_RM_ and cross-reactive HA-B_RM_ against H3N2 A/Uruguay/716/07 strain (Urg) ^14^ using flow cytometry. T_RH_ ablation resulted in diminished strain-specific (X31 HA^+^ and Urg HA^-^) and cross-reactive (both X31 HA^+^ and Urg HA^+^) B_RM_ responses (Fig. 7k), indicating potential roles of T_RH_ cells in the development of both strain-specific and cross-reactive B cell immunity.

Since CD8^+^ T_RM_ (particularly NP_366-374_ T_RM_ ^40^) and possibly B_RM_ cells are important in mediating host heterologous protection ^5^, we next sought to determine whether the ablation of T_RH_ cells impairs host protective immunity against heterologous virus infection. To do so, we employed a heterologous infection and challenge model in which X31 virus was used as the primary infection and lethal PR8 virus was used as secondary challenge ^55^. PR8 and X31 viruses differ in the viral surface proteins but share internal viral proteins such as NP ^64^. As such, CD8^+^ T_RM_ and possibly B_RM_ cells against internal viral epitopes (mainly against viral NP protein) can provide heterologous protection ^5, 40^. We infected WT or *Bcl6*^*ΔCD4ERT2*^ mice with X31 virus and treated the mice with tamoxifen. We confirmed that T_RH_ ablation affects CD8 NP_366-374_ T_RM_, B_GC_ and B_RM_ development in the X31 model at 35 d.p.i. (Extended Fig. 7f). We then re-challenged the mice with a lethal dose of PR8 in the presence of FTY720 to block the contribution of circulating memory CD8^+^ T and B cells at 42 d.p.i. (Fig. 7l). Close to 40% of *Bcl6*^*ΔCD4ERT2*^ mice succumbed, while WT mice were fully protected with lethal PR8 infection (Fig. 7m). Thus, we conclude that T_RH_ cells are required for the optimal protection against secondary heterologous infection, most likely through their help for the development of protective CD8 T_RM_ and B_RM_ response.

### Identification of the factors mediating T_RH_-derived “local” help

We next sought to identify the underlying mechanisms by which T_RH_ cells promote B cell and CD8^+^ T_RM_ immunity. As shown in Fig. 2, T_RH_ cells expressed high levels of *Il21*. To identify the major IL-21-expressing cell types in the lung, IL-21-VFP reporter mice were infected with PR8 and examined at 28 d.p.i. The vast majority of lung IL-21-VFP^+^ cells were CD4^+^ T cells (Fig. 8a). IL-21-VFP^hi^ cells, which expressed the highest levels of IL-21 than those IL-21-VFP^low^ cells (Extended Fig. 8a), were mainly T_RH_ cells (Fig. 8b). Additionally, T cell-specific BCL6 deficiency impaired *Il21* expression in the lungs (Extended Fig. 8b). These data suggest that T_RH_ cells are the main IL-21 producers in the lungs following influenza virus infection.

**Fig. 8.**
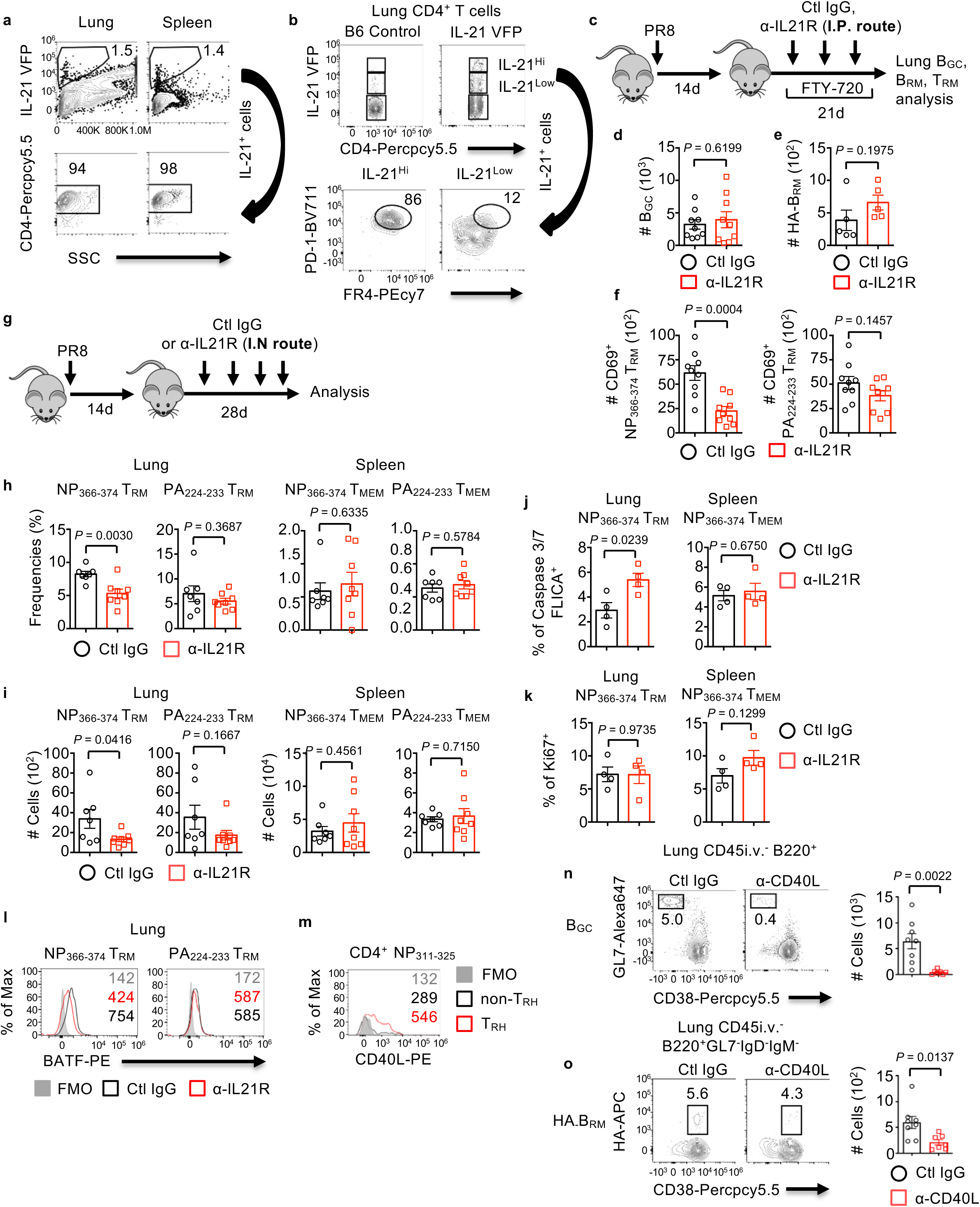
IL-21 or CD40L-dependent T_RH_ help to CD8^+^ or B cells. **a-b**, IL-21-VFP reporter mice were infected with PR8. **a**, IL-21-VFP expressing cells in the lungs or spleens were identified by flow cytometry at 28 d.p.i. **b**, Representative dot plot of IL-21^Hi^ or IL-21^Low^ CD4^+^ T cells that were PD-1^Hi^FR4^Hi^. **c-f**, WT mice were infected PR8 with or without IL-21R blockade through intraperitoneal (I.P.) route starting at 14 d.p.i. in the presence of FTY-720 administration (13-34 d.p.i.). **c**, Experimental scheme. Cell numbers of lung parenchymal B_GC_ (**d**), HA-specific B_RM_ (**e**) and CD8^+^CD69^+^ NP_366-374_ or CD8^+^CD69^+^ PA_224-233_ T_RM_ cells (**f**). **g-k**, WT mice were infected with PR8 with or without IL-21R blockade through intranasal (I.N.) route at 14 d.p.i. **g**, Experimental scheme. **h-i**, Frequencies (**h**) or cell numbers (**i**) of lung tissue CD8^+^CD69^+^ NP_366-374,_ CD8^+^CD69^+^ PA_224-233_ T_RM_, splenic CD8^+^ NP_366-374_ or PA_224-_233 memory T cells (T_MEM_) at 42 d.p.i. **j**, Percentage of apoptotic cells were identified by active caspase 3/7-FLICA staining within lung CD8^+^ NP_366-374_ T_RM_ or splenic CD8^+^ NP_366-374_ T_MEM_ at 28 d.p.i. **k**, percentages of proliferating cells were identified by Ki67 staining within lung CD8^+^ NP_366-374_ T_RM_ or splenic CD8^+^ NP_366-374_ T_MEM_ at 28 d.p.i. **l**, Representative histogram of BATF expression in lung CD8^+^ NP_366-374_ or CD8^+^ PA_224-233_ T_RM_ of mice received with ctl IgG or α-IL21R at 35 d.p.i. **m**, Representative histogram of CD40L expression in influenza-specific lung T_RH_ or non-T_RH_ at 28 d.p.i. **n-o**, WT mice were infected with PR8 and received ctl IgG or α-CD40L weekly (I.P. route) starting at 14 d.p.i. in the presence of daily FTY-720 administration (13-34 d.p.i.). Representative dot plot or cell numbers B_GC_ (**n**) and HA-specific B_RM_ (**o**) cells at 35 d.p.i. In **a-b and j-m**, representative data were from at least two independent experiments (4-5 mice per group). In **d, f-i** and **n-o**, data were pooled from two independent experiments (3-5 mice per group). *P* values of all experiments were calculated by unpaired two-tailed Student’s t-test.

Since IL-21 is an important cytokine that has been implicated in facilitating B_GC_, memory B and CD8^+^ effector and memory T cell responses ^65^, we blocked IL-21 signaling starting at 14 d.p.i. through intraperitoneal injection of α-IL-21R in the presence of FTY720 (Fig. 8c). To our surprise, IL-21 signaling blockade did not impair B_GC_ or influenza HA-specific B_RM_ responses in the lung tissue (Fig. 8d, e). However, IL-21 signaling blockade significantly diminished CD8^+^ NP_366-374_ T_RM_ responses, suggesting that T_RH_ cells help CD8^+^ T_RM_ establishment and/or maintenance through IL-21 (Fig. 8f). We also blocked IL-21 signaling locally in the lung through intranasal delivery of α-IL-21R (Fig. 8g). Lung local blockade of IL-21 signaling diminished lung CD8^+^ NP_366-374_ T_RM_ magnitude, but not splenic memory T cells (Fig. 8h, i). Local blockade of IL-21 signaling also did not affect B_GC_ or HA-specific B cell responses in the lung parenchymal compartment (Extended Fig. 8c).

Consistent with the diminished CD8^+^ NP_366-374_ T_RM_ responses, IL-21R signaling blockade leaded to enhanced cellular apoptosis but not proliferation, specifically in CD8^+^ NP_366-374_ T_RM_ but not those of splenic memory T cells (Fig. 8j, k). Of note, compared to CD8^+^ PA_224-233_ T_RM_ cells, CD8^+^ NP_366-374_ T_RM_ cells exhibited higher expression of genes associated with IL-21 signaling including *Batf* (Extended Fig. 8d, e) ^66^. These data suggest that CD8^+^ NP_366-374_ T_RM_ cells, but not CD8^+^ PA_224-233_ T_RM_ cells, potentially receive IL-21 signaling in the lungs at the memory stage following influenza virus infection. Consistent with the data, we found that IL-21R blockade diminished BATF expression in CD8^+^ NP_366-374_ T_RM_, but not CD8^+^ PA_224-233_ T_RM_ cells at 35 d.p.i. (Fig. 8l). Taken together, these data suggest T_RH_ cells provide local help for the development of CD8^+^ NP_366-374_ T_RM_ cells in an IL-21-dependent manner.

CD40-CD40L interaction is critical for T cell help of B_GC_ cell responses ^5, 57^. Previously, it was shown that lung B_GC_ cells contributed to lung local B_RM_ cell responses ^14^. Furthermore, diminished B_GC_ response following CD40L blockade can lead to impaired B_RM_ formation when α-CD40L was given before 20 d.p.i., although it is unknown whether this is due to diminished lung or circulating B_GC_ responses ^5^. Lung influenza-specific T_RH_ cells express higher levels of CD40L than non-T_RH_ cells (Fig. 8m and Extended Fig. 8f). To determine whether CD40L promotes lung B_GC_ responses to facilitate local B_RM_ formation following influenza infection, we inoculated α-CD40L into PR8-infected mice at 14 d.p.i. in the presence of FTY720 as of Fig. 8c. As shown in Fig. 8n and o, CD40L blockade greatly diminished lung B_GC_ formation and the magnitude of HA-specific B_RM_ responses in the lung tissue. These data indicate that T_RH_ cells facilitate B_GC_ and B_RM_ responses through CD40L-dependent mechanisms.

## Discussion

In this report, we have discovered a previously unrecognized requirement for “*in situ*” CD4^+^ T cell help in the respiratory mucosa, which is mediated by what we have termed T_RH_ cells, in the development of localized protective memory responses following influenza virus infection. T_RH_ cells co-manifest phenotypic and transcriptional hallmarks of both T_FH_ and T_RM_ cells. We have further identified the cellular and molecular mechanisms guiding the development of T_RH_ cells, and key factors mediating their helper function to B and CD8^+^ T cells (Extended Fig. 8g).

The expression of T_FH_-associated molecules by T_RH_ cells appear to be lower compared to those of splenic T_FH_ cells (such as BCL6 and CXCR5). These PD-1^Hi^ CXCR5^Low^ BCL6^Int^ T_FH_-like cells have been previously observed ^34, 35, 36, 37, 38^. However, the cellular identity and developmental cues regulating their development remain largely elusive. Furthermore, the physiological function of these cells beyond their help on the generation of B_GC_ cells are currently unknown. Using total and single cell transcriptional profiling and phenotypic analysis combined with parabiosis, we have identified that these tissue T_FH_-like cells exhibit enhanced expression of molecules-associated with peripheral migration and tissue residency, and show limited circulating ability. We therefore term these cells as T_RH_ cells based on their transcriptional signature, non-migratory features and helper function. Currently, the cellular origins of these T_RH_ cells are not determined in this study. T cell priming occur in the draining lymph nodes and it is possible that T_RH_ cells originate from those lymph node pre-T_FH_ or interfollicular T_FH_ cells (T_FH_ precursors outside the GC ^67, 68^) entering circulation and adopting T_RM_ signatures following entry in the lung parenchymal environment. Conversely, it is also possible that T_RH_ cells develop from Th1-polarized cells and adopt “T_FH_-like” features in the iBALT structure when interacting with B cells. These possibilities of T_RH_ cell development warrant further investigation. Nevertheless, based on the T_RH_ transcriptional profiles and the dual requirement of BCL6 and Bhleh40 for their formation, we believe that T_RH_ cells are likely a “hybrid” of T_FH_ and T_RM_ cells, and may represent a unique population present in various tertiary lymphoid structures of mucosal tissues.

Lung T_RH_ cells appear to be required for programing localized memory B and T cell responses in the respiratory mucosal tissue. Previously, it was shown that locally-generated lung B_GC_ cells contribute significantly to the lung B_RM_ cell pool following influenza virus infection ^14^. Thus, T_RH_ cells assist the development of B_RM_ responses likely due to their help for the generation of lung B_GC_ cells, rather than their direct “help” on lung B_RM_ cells per se for their differentiation and/or maintenance. Consistent with this idea, the ablation of T_RH_ cells around 4 weeks post infection did not significantly impact lung B_RM_ cell maintenance. In accordance, the inoculation of CD40L Ab after 3 weeks of infection, which blocked the generation of lung B_GC_ cells, fails to impact B_RM_ cell generation ^5^. These data suggest that localized B_GC_ responses in the iBALT, programmed by lung T_RH_ cells in the first three weeks of infection, facilitate B_RM_ development following influenza virus infection.

Beyond their help to B cells, T_RH_ cells are required for the optimal responses of a protective CD8^+^ T_RM_ population, most likely through the production of IL-21. Of note, it was previously shown that continuous CD4^+^ T cell help beyond T cell priming was not required for the differentiation of CD103^+^ CD8^+^ T_RM_ cells ^30^. Consistently, we found that CD4^+^ T cell depletion following the resolution of primary infection did not impact CD103 expression by CD8^+^ T_RM_ cells. However, persistent “late” help following viral clearance is required for the generation and/or the maintenance of a protective lung T_RM_ population in our study. The timing of CD4^+^ T cell depletion (day 7 previously ^30^ versus day 14 CD4^+^ T cell depletion used here) likely explains the differences observed in the two models. It is possible that early depletion of CD4^+^ T cells before viral clearance (i.e. day 7) alters lung viral and/or inflammatory environment, which compensate the requirement of lung CD4^+^ T cells for CD8^+^ T_RM_ programming and/or maintenance. Other possibilities, including infection schemes and the levels of CD4^+^ T cell depletion following α-CD4 inoculation may also contribute to the differences observed. Nevertheless, using multiple lines of approaches including high or low doses of α-CD4 depletion, inducible CD4^+^ T cell-specific BCL6 ablation and IL-21 blockade combined with long-term FTY720 treatment, we have provided comprehensive evidence that localized CD4^+^ T cell help, mediated by T_RH_ cells and IL-21, is required for the optimal generation of a population of protective T_RM_ cells (H2D^b^-restricted NP_366-374_ T_RM_ cells) following the resolution of primary infection.

Then, the question is why lung T_RH_ help, in the form of IL-21 provision, is specifically required for CD8^+^ NP_366-374_ T_RM_ responses. We have shown previously that CD8^+^ NP_366-374_ T_RM_ cells receive constant antigen engagement in the lungs at the memory stage due to the chronic deposition of high levels of influenza NP antigen ^40^. Compared to conventional CD8^+^ PA_224-233_ T_RM_ cells, CD8^+^ NP_366-374_ T_RM_ cells express high levels of PD-1 and exhibit “exhausted-like” phenotypes similar to those of T cells receiving chronic antigen exposure during chronic viral infections ^40, 69, 70^. Consequently, the maintenance of these T_RM_ cells requires persistent MHC I-dependent stimulation at the memory stage ^40^, similar to those of exhausted CD8^+^ T cells ^69^. Notably, CD4^+^ T cell help in the form of IL-21 has been recently demonstrated to sustain those PD-1-expressing CD8^+^ T cells during chronic viral infection and tumor growth ^71^. Thus, CD4^+^ T cell help and IL-21 signaling may be specifically required for maintaining the survival of lung CD8^+^ T_RM_ cells receiving persistent low levels of *in situ* antigenic stimulation. Consistent with this idea, NP_366-374_ T_RM_ cells express higher levels of molecules associated with IL-21 signaling, particularly BATF.

Influenza virus is able to undergo antigenic shift, drift and re-assortment to escape previously established host immunity. Current influenza vaccines require yearly update and only provide high levels of protection when influenza vaccine strains match exactly with the circulating strains. Much of attention has been given to the development of potential universal influenza vaccines recently. The common goals of various universal influenza vaccine candidates are to induce broadly neutralizing influenza Ab, strong CD8^+^ memory T cell responses against conserved epitopes and/or high levels of localized lung-resident mucosal immunity that can restrict viral spreading early ^72, 73, 74, 75^. Due to the high mutation rates of influenza viruses, it is conceivable that the induction of “all inclusive” immune responses, i.e. induction of strong both B and CD8^+^ T cell immunity in the mucosal tissue, may be required for the ultimate success of a universal influenza vaccine ^75^. Due to the unique ability of T_RH_ cells in assisting local B_RM_ and T_RM_ development, it is tempting to speculate that the specific promotion of T_RH_ cells may simultaneously promote both memory B and CD8^+^ T cell immunity in the respiratory tract during vaccination, thereby providing rapid cross-reactive protection against broad-spectrum viruses.

Acute influenza virus infection can lead to the development of chronic lung pathogenic sequelae 40, 76. The roles of iBALT and T_RH_ in the development of chronic lung conditions following influenza infection needs further investigations. Additionally, tertiary lymphoid (B/T) aggregates or iBALT-like structures have been observed in many chronic lung diseases including asthma, pulmonary fibrosis, COPD (Chronic obstructive pulmonary disease) etc ^77, 78, 79, 80^. It was speculated that these tertiary lymphoid structures may play important roles in modulating the disease progression in those chronic lung conditions ^47, 79^. T_RH_ cells may therefore participate in the regulation of chronic lung disease development. Further, recent advances have suggested that the development of tertiary lymphoid structures consisting of CD4^+^ T and B cell clusters inside tumor effectively predicts patient responses to checkpoint blockade (anti-PD-1 and/or anti-CTLA4) therapies ^81, 82, 83^. Given the transcriptional similarity of T_RM_ cells and tumor infiltrating lymphocytes (TILs) ^84^, it is possible that those T_FH_-like cells present inside those tertiary lymphoid structures within tumor also exhibit tissue-resident signatures (i.e. T_RH_-like cells). Indeed, a previous report has suggested the development of CXCL13^+^ (human T_FH_ marker) BHLHE40^+^ CD4^+^ T cells are associated with enhanced responsibility to anti-PD therapy in colorectal cancer ^85^. Thus, it is possible that T_RH_-like cells may also provide “*in situ*” help for the optimal generation and/or maintenance of anti-tumor CD8 TILs following checkpoint blockade.

In summary, our data have revealed a complex T-B interaction network that is programed by lung T_RH_ cells for the maintenance of protective local respiratory immunity following acute influenza virus infection. Moving forward, it is of substantial interest to dissect the mechanisms modulating the development of T_RH_ cells and the precise function of T_RH_ cells in assisting the development of local B and CD8 T cell immunity during infection, vaccination and possibly cancer.

## Materials and methods

### Mice and influenza viral infection

WT C57BL/6, CD45.1 and IL-21 VFP reporter mice were purchased from the Jackson Laboratory (JAX) and bred in-house. To generate CD45.1^+^ and CD45.2^+^ (CD45.1^+^/CD45.2^+^) mice, CD45.1 mice were crossed with C57BL/6 mice. *Bcl6*^*fl/fl*^ were generated as previously reported ^63^. *Bcl6*^*ΔT*^ were generated by crossing with CD4-Cre transgenic. *Bcl6*^*ΔCD4ERT2*^ were generated by crossing with CD4-ERT2 transgenic mice. *Bcl6*^*fl/fl*^ or *Bcl6*^*ΔCD4ERT2*^ mice were additionally crossed with Rosa26 LSL-YFP (JAX) mice for the determination of the efficiency of tamoxifen induced gene recombination. *Bhlhe40*^*fl/fl*^ or *Bhlhe40*^*ΔT*^ mice were generated as previously reported ^55^. All animal protocols were approved by the Institutional Animal Care and Use Committees (IACUC) of the Mayo Clinic (Rochester, MN). Sex-matched and age-matched 8-to 10-week-old mice of both sexes were used in the experiments. Influenza A/PR8/34 (∼200 pfu/mouse in the primary infection and ∼1×10^4^ pfu/mouse in the secondary infection) and Influenza A X31 (∼800 pfu/mouse in the primary infection) were infected into the mice by intranasal (i.n.) under anesthesia as reported before ^42^.

### Intravascular labeling with α-CD45 and preparation of lung cell suspension

Mice were intravenously (i.v.) injected with 2 µg of **α**-CD45 (Clone: 30-F11) (Tonbo Biosciences) diluted in 300 µL of sterile PBS five minutes before sacrifice. To prepare single cells from lung tissue, lungs were cut into small pieces, digested with Type 2 Collagenase (Worthington Biochemical) and dissociated in 37°C for 30 min with Gentle-MACS (Miltenyi). Cells were further ground through 70µm cell strainer (Falcon) and washed with plain IMDM (Gibco). After red blood cell lysis, cells were centrifuged and re-suspended in cold FACS buffer (PBS, 2 mM EDTA, 2 % FBS, 0.09 % Sodium Azide) for flow cytometry analysis. Lung circulating immune cells are i.v. Ab^+^ and lung tissue immune cells are defined by i.v. Ab^-^. Lung T_RM_ cells were defined as CD45_i.v._^-^CD8^+^ tetramer^+^CD69^+^. Lung B_RM_ cells were defined as CD45_i.v._^-^ B220^+^ GL7^-^ IgD^-^ IgM^-^ CD38^+^ influenza B cell antigen (HA or NP)^+^.

### Antibody administration *in vivo*

Influenza infected WT mice were administrated with control IgG or various neutralizing or depleting Ab as described in the text. For CD4 T cell depletion with high dose of α-CD4, mice were injected with 250 µg α-CD4 weekly (Clone: GK1.5, BioXCell) starting at 14 d.p.i. For circulating CD4 T cell depletion, mice were I.P. injected with 40 µg α-CD4 for the first dose followed with 10 µg α-CD4 weekly. For B cell depletion, mice were injected with 500 µg of α-CD20 (Clone: 5D2, Genentech). CD40L blockade or iBALT depletion were achieved by the injection of 250 µg of α-CD40L (Clone: MR-1, BioXCell) or 250 µg of LTβR-Ig weekly respectively. For systemic IL-21R blockade, 500 µg of α-IL21R (Clone: 4A9, BioXCell) was injected through I.P weekly starting at 14 d.p.i. For lung local IL-21R blockade, 50 µg of α-IL21R was injected through intranasal route (I.N.) weekly starting at 14 d.p.i. In some experiments, FTY720 (1 mg/kg) (Cayman) was administrated daily from 13 d.p.i. to block lymphocyte migration until mouse euthanasia.

### Tamoxifen treatment

To induce gene recombination in *Bcl6*^*ΔCD4ERT2*^ mice, tamoxifen (Sigma-Aldrich) was diluted in warm sunflower oil (Sigma-Aldrich) and daily treated via intraperitoneal route for 5 consecutive times. Each application was 2 mg per mouse.

### Flow cytometry analysis

For cell surface staining, cells were incubated with the appropriate antibody cocktail with FACS buffer for 30 min at 4 °C dark condition. Then cells were washed with FACS buffer. For intracellular staining, cell suspensions were stained with indicated surface markers and then washed with FACS buffer. Cells were then fixed and permeabilized with either Perm Fix and Wash buffer (Biolegend, for cytokine staining) or the Foxp3 transcription factor staining buffer set (eBioscience, for KI-67, Foxp3, BATF and BCL6 staining) for 1 hour at room temperature (RT) in the dark. Cells were washed twice with Perm Wash buffer (Biolegend or eBioscience) and stained with indicating Abs for 1 hour at RT. After staining, cells were washed again with Perm Wash buffer before flow cytometry acquisition. FACS Abs were primarily purchased from Biolegend, BD Biosciences, eBioscience or Tonbo Biosciences. The clone number of those Abs are as follows: CD45 (Clone: 30-F11), CD45.1 (Clone: A20), CD45.2 (Clone: 104), CD4 (Clone: RM4-5), CD44 (Clone: IM7), PD-1 (Clone: 29F.1A12), FR4 (Clone: eBio12A5), GITR (Clone: DTA-1), B220 (Clone: RA3-6B2), FAS (Clone: SA367H8), GL7 (Clone: GL7), CD38 (Clone: 90), IgD (Clone: 11-26c.2a), IgM (Clone: 11/41), CD8a (Clone: 53-6.7), CD69 (Clone: H1.2F3), CD103 (Clone: 2E7), CXCR5 (Clone: SPRCL5), ICOS (Clone: 7E.17G9), P2RX7 (Clone: 1F11), CXCR6 (Clone: SA051D1), CD40L (Clone: MR1), BCL6 (Clone: K112-91), Foxp3 (Clone: 3G3), BATF (Clone: S39-1060), Ki67 (Clone: SoLA15), Streptavidin, IFN-γ (Clone: XMG1.2), IL-17 (Clone: TC11-18H10.1) and IL-4 (Clone: 11B11). The expression of Bhlhe40 was measured with Primeflow kit (Thermo Fisher Scientific). For CD40L staining, lung cells were pre-activated with 100 ng/mL of PMA and 1 µg/mL of ionomycin (Sigma-Aldrich) for 3 hrs in the 37°C before CD40L surface staining. H-2D^b^-NP_366–374_, H-2D^b^-PA_224–233_ and I-A^b^-NP_311-325_ tetramers were obtained from the National Institutes of Health tetramer facility. After Ab staining, cells were acquired with an Attune NxT system (Life Technologies). Data analysis was performed by FlowJo software (Tree Star).

### B cell antigens

Influenza PR8-HA protein was a gift from Dr. Michelle C. Crank (NIH). PR8-NP protein was purchased from Sino Biological. Purified antigens were biotinylated using an EZ-Link Sulfo-NHS-LCBiotinylation kit (Thermo Fisher Scientific) using a 1:1.3 M ratio of biotin to Ag. To make tetramers, biotinylated Ags were mixed with streptavidin–PE (PJ27S; ProZyme) at the ratio determined or at a 5 to 1 ratio using the biotin concentration provided by the manufacturer as described before ^86^. Following a 30-min incubation on ice, unconjugated biotinylated Ag was often removed by several rounds of dilution and concentration using a 100 kDa Amicon Ultra (MilliporeSigma) or 300 kDa Nanosep centrifugal devices (Pall). Tetramers were stored at 1 µM in 1× DPBS at 4°C prior to use. H3N2 X31-HA conjugated with APC and H3N2 Urg (Uruguay)- HA conjugated with PE were reported before ^14^.

### Apoptosis analysis

Apoptosis of NP_311-325_ T_RH_, non-T_RH_ cells or lung NP_366-374_ T_RM_, splenic NP_366-374_ T_MEM_ were assessed with CellEvent™ Caspase-3/7 Green Flow Cytometry Assay kit (ThermoFisher). Lung single cells were stained with surface markers then incubated with caspase 3/7 green detection reagent for 30 minutes at 37°C as described in the manufacturer’s instructions. Casepase 3/7 Flica activities were analyzed flow cytometry.

### Immunohistochemistry and immunofluorescence

Left lobe of whole lungs was harvested and fixed in 10% formaldehyde (Fisher Scientific) until embedding. Fixed lung tissues were embedded in paraffin, sectioned at 10-µm thickness. To identify tertial lymphoid structure, the lung tissue slide was stained with hematoxylin and eosin by the Mayo Clinic Histology Core Laboratory (Scottsdale, AZ). To measured inducible bronchus-associated lymphoid tissues (iBALT) structure, lung tissue sections were deparaffinized in CitriSolv (Fisher Scientific) for 30 min and then immersed in ethanol series from 100, 95, 85, and 75% to distill H_2_O for 5 min each for tissue hydration. For antigen retrieval, hydrated sections were steamed for 20 min in 1 mM EDTA (pH 8.0). For detecting, IL-21 expressing CD4^+^ T cells in iBALT, left lobe of lungs from influenza infected-IL21-VFP reporter mice were harvested and fixed for overnight at room temperature in 4% paraformaldehyde (PFA). The tissues were incubated in 10% sucrose for overnight, then incubated in 30% sucrose before being embedded in OCT compound and cryosectioned. The frozen sections were fixed with cold acetone for 20 min. Lung sections were blocked with Super Block Blocking buffer (Fisher Scientific) for 1 hr at RT. Anti-B220-eflour660 (Clone: 4SM95, Invitrogen), -CD4-eflour570 (Clone: RA3-6B2, Invitrogen), -GL7-Alexa488 (Clone: GL7, Biolegend) and/or -GFP-Alexa488 (Clone: FM264G) were stained on the lung tissue sections for overnight at 4°C. After washing in 0.1% PBST (PBS with Tween 20), the slides were counterstained with 4’, 6-diaminodino-2-phenylindole (DAPI) and mounted. Tissue staining was reviewed and representative images were acquired on a Zeiss LSM 780 confocal system (Carl Zeiss).

### Quantitative RT-PCR

Total RNA was extracted from lung tissue or sorted cells as indicated in the text with Total RNA purification kit (Sigma) and treated with DNase I (Invitrogen). Random primers (Invitrogen) and MMLV reverse transcriptase (Invitrogen) were used to synthesize first-strand cDNAs from equivalent amounts of RNA from each sample. These cDNA was subjected to realtime-PCR with Fast SYBR Green PCR Master Mix (Applied Biosystems). Realtime-PCR was conducted in duplicates in QuantStudio3 (AppliedBioscience). Data were generated with the comparative threshold cycle (Delta CT) method by normalizing to hypoxanthine phosphoribosyltransferase (HPRT). Sequences of primers used in the studies are listed as following. *Hprt*-F: CTCCGCCGGCTTCCTCCTCA, *Hprt*-R: ACCTGGTTCATCATCGCTAATC. *Il21*-F: CGCTCACGAATGCAGGAGTA, *Il21*-R: GTCTGTGCAGGGAACCACAA.

### Parabiosis surgery

To examine tissue residency of lung T_RH_ or non-T_RH_ cells, parabiotic surgery was performed. CD45.1^+^ or CD45.1^+^/CD45.2^+^ mice were infected with influenza PR8. 3 weeks later, mice were anesthetized with ketamine and xylazine and shaved in lateral skin area. After disinfection, shaved skin area was made an incision then matched from the olecranon to the knee joint of each mouse. Matching area was opposed with continuous sutures. Parabionts were then allowed to rest for 14 d before analysis. Equilibration of parabionts was confirmed in the peripheral blood before tissue analysis.

### Nanostring analysis

To perform nanostring analysis, influenza-specific lung CD8 T_RM_, splenic T_MEM_, lung T_RH_, lung non-T_RH_ and splenic T_FH_ cells were sorted as indicated in the text. Total RNA was extracted from sorted T cell populations with mini RNA-easy Kit (Qiagen). Equal amounts of total RNA from different groups were used for the assay. Hybridization reaction was established by following the instruction of the manufacturer. Aliquots of Reporter CodeSet and Capture ProbeSet were thawed at RT. Then, a master mix was created by adding 70 µl of hybridization buffer to the tube containing the reporter codeset. Eight microliters of this master mix was added to each of the tubes for different samples; 5 µl (50 ng) of the total RNA sample was added into each tube. Then, 2 µl of the well-mixed Capture probeset was added to each tube and placed in the preheated 65°C thermal cycler. All the sample mixes were incubated for 16 hours at 65°C for completion of hybridization. The samples were then loaded into the sample hole in the cartridge and loaded into the NanoString nCounter SPRINT Profiler machine (NanoString). When the corresponding Reporter Library File (RLF) running is finished, the raw data were downloaded and analyzed with NanoString Software nSolver 3.0 (NanoString). mRNA counts were processed to account for hybridization efficiency, background noise, and sample content, and were normalized using the geometric mean of housekeeping genes. All data were normalized by housekeeping genes. Heat map was generated by MeV software.

### Single-cell RNA sequencing

Sorted CD45_i.v._^-^CD4^+^CD44^Hi^ T cells from pooled lung cells of mice (5 mice) infected with influenza virus (28 d.p.i.) were loaded on the Chromium Controller (10x Genomics). Single-cell libraries were prepared using the Chromium Single Cell 3’ Reagent kit (10x Genomics) following manufacturer’s instruction. Paired-end sequencing was performed using an Illumina HiSeq 2500 in rapid-run mode. CellRanger software package (10x Genomics) were used to align and quantify sequencing data from 10x Genomics. All scRNA-seq analyses were performed in R using the package Seurat (version 2.0) ^87^.

### Total RNA-sequencing

Lung CD45_i.v._^-^CD4^+^CD44^Hi^GITR^-^PD-1^Hi^FR4^Hi^, CD45_i.v._^-^CD4^+^CD44^Hi^GITR^-^PD-1^Low^FR4^Low^, splenic CD4^+^CD44^Hi^GITR^-^PD-1^Hi^ CXCR5^+^ T_FH_ or splenic CD4^+^CD44^Hi^GITR^-^PD-1^low^ CXCR5^-^non-T_FH_ cells were sorted from total of 15 mice that were infected with influenza virus (28 d.p.i.). RNA was extracted using RNeasy Plus Mini Kit (Qiagen) following the manufacture’s recommendation. After quality control, high quality (Agilent Bioanalyzer RIN >7.0) total RNA was used to generate the RNA sequencing library. cDNA synthesis, end-repair, A-base addition, and ligation of the Illumina indexed adapters were performed according to the TruSeq RNA Sample Prep Kit v2 (Illumina, San Diego, CA). The concentration and size distribution of the completed libraries was determined using an Agilent Bioanalyzer DNA 1000 chip (Santa Clara, CA) and Qubit fluorometry (Invitrogen, Carlsbad, CA). Paired-end libraries were sequenced on an Illumina HiSeq 4000 following Illumina’s standard protocol using the Illumina cBot and HiSeq 3000/4000 PE Cluster Kit. Base-calling was performed using Illumina’s RTA software (version 2.5.2). Paired-end RNA-seq reads were aligned to the mouse reference genome (GRCm38/mm10) using RNA-seq spliced read mapper Tophat2 (v2.1.1). Pre- and post-alignment quality controls, gene level raw read count and normalized read count (i.e. FPKM) were performed using RSeQC package (v2.3.6) with NCBI mouse RefSeq gene model. For functional analysis, GSEA was used to identify enriched gene sets, from the hallmark collection of MSigDB, having up-regulated and down-regulated genes, using a weighted enrichment statistic and a log2 ratio metric for ranking genes.

### Quantification and Statistical Analysis

Statistical analysis was done using GraphPad Prism 7.0 (GraphPad Software) and presented as means ± SEM. Unpaired or paired Student t tests and one-way or two-way ANOVA analysis were used in data analysis. Logrank test was used for analysis of survival study.

## Acknowledgements

We thank NIH tetramer core for tetramers and Mayo flow cytometry core for technical assistance. We thank Genetech for α-CD20 and Drs. Barney Graham and Michelle Crank for PR8 HA protein used in this study. This study was funded by the US National Institutes of Health RO1s AI112844, AI147394 and AG047156 (J.S.), R01 AI125741 and RO1 AI148403 (W.C.) and R01 AI057459 (M.H.K.).

## Figure Legends

**Extended Fig. 1.**
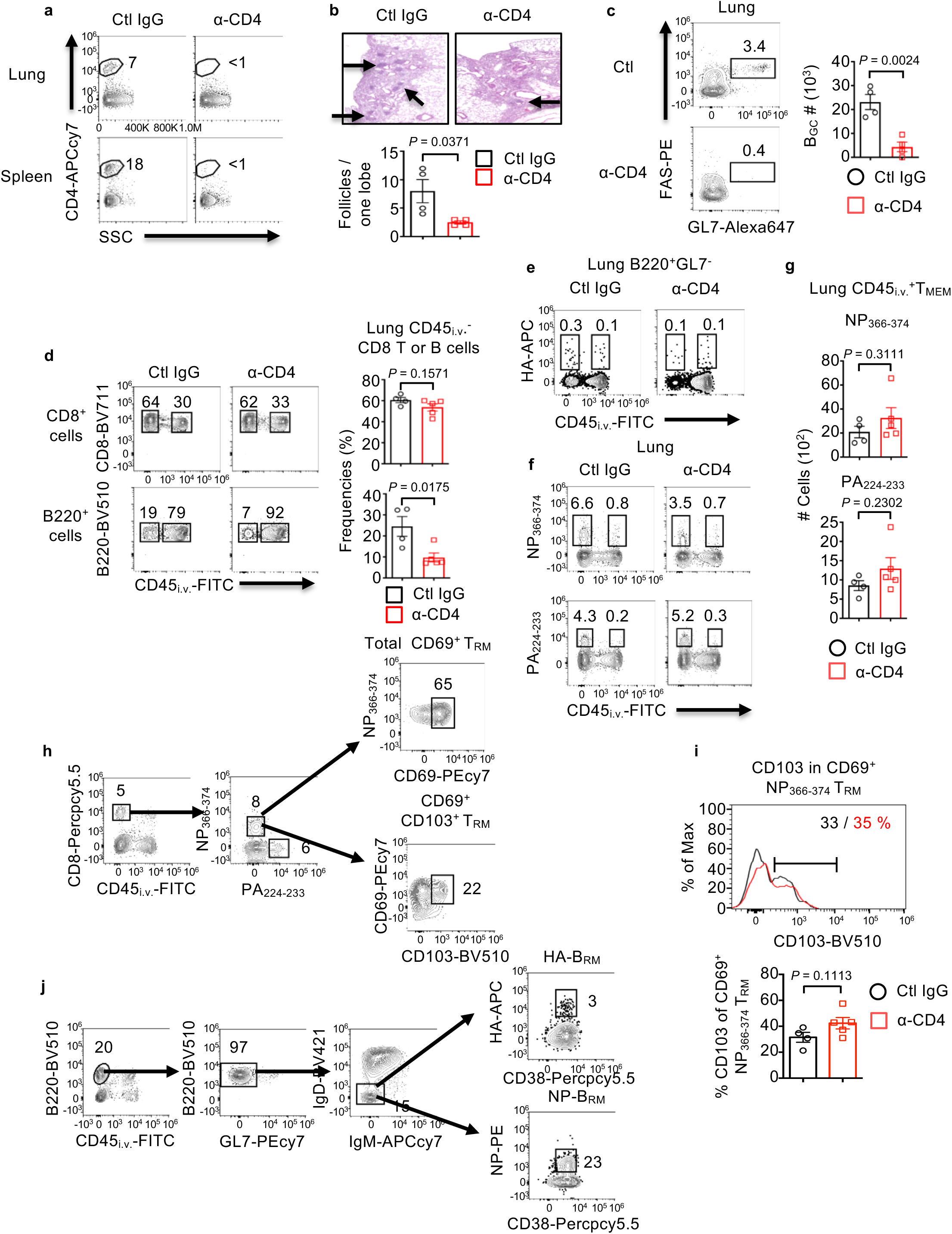
“Late” CD4^+^ T cell help shapes respiratory mucosal memory CD8^+^ and B cell immunity. WT mice were infected with PR8 and treated with ctl IgG or α-CD4. **a**, The efficiency of CD4^+^ T cell depletion in the lung or spleen. **b**, Representative image of lung section stained with H&E. Black arrows indicate tertial lymphoid structure. **c**, Representative dot plot and cell numbers of lung germinal center B (B_GC_) cells. **d**, Frequencies of lung circulating (CD45_i.v._^+^) or parenchymal (CD45_i.v._^-^) CD8^+^ T or B cells in mice treated with control IgG or α-CD4. **e**, Influenza HA-specific B cells (HA-B) in the lungs were identified through HA antigen staining. **f**, Lung tissue or circulating CD8^+^ NP_366-374_ or PA_224-233_ T cells were identified through CD45_i.v._ staining and analyzed by flow cytometry. **g**, Lung circulating CD8^+^ NP_366-374_ or CD8^+^ PA_224-233_ T cells were enumerated. **i**, Histogram of CD103 expression or frequency of CD103^+^ cells within CD8^+^CD69^+^ NP_366-374_ T_RM_ cells. **h, j**, Gating strategies of CD8^+^ T_RM_ (**h**) or B_RM_ (**j**) cells in the lung. Representative data were from at least two independent experiments (4-5 mice per group). *P* values were calculated by unpaired two-tailed Student’s t-test.

**Extended Fig. 2.**
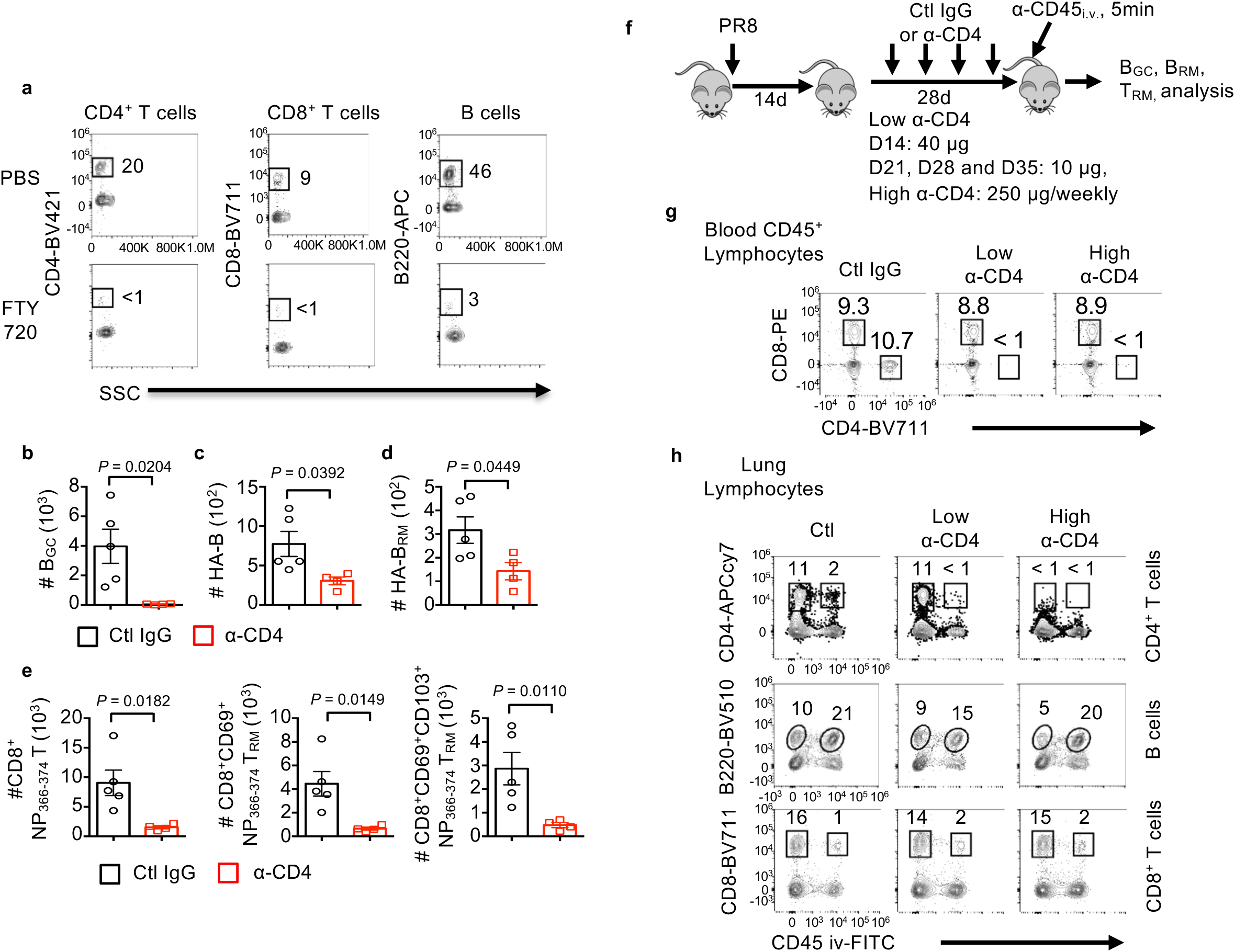
Lung “local” CD4 T cell help for the development robust memory CD8^+^ and B cell immunity. WT mice were infected with influenza PR8 (**a**) or X31 (**b-e**) and treated with control IgG or α-CD4 (starting at 14 d.p.i.) in the presence of daily FTY720 (starting at 13 d.p.i.). **a**, Blood lymphocytes in the PBS or FTY720 administrated mice. **b-e**, Numbers of lung B_GC_ cells (**b**), total HA specific B cells (**c**), HA-specific B_RM_ (**d**), and total CD8^+^ NP_366-374_ memory T cells, CD8^+^ CD69^+^ NP_366-374_ T_RM_ or CD8^+^ CD69^+^ CD103^+^ NP_366-374_ T_RM_ cells (**e). f-h**, WT mice were infected with PR8 and received with ctl IgG, low or high dose of α-CD4 (starting at 14 d.p.i.). Experimental scheme (**f**), representative dot plot of blood lymphocyte population (**g**) and lung lymphocyte population (**h**). Representative data were from at least two independent experiments (4-5 mice per group). *P* values were calculated by unpaired two-tailed Student’s t-test.

**Extended Fig. 3.**
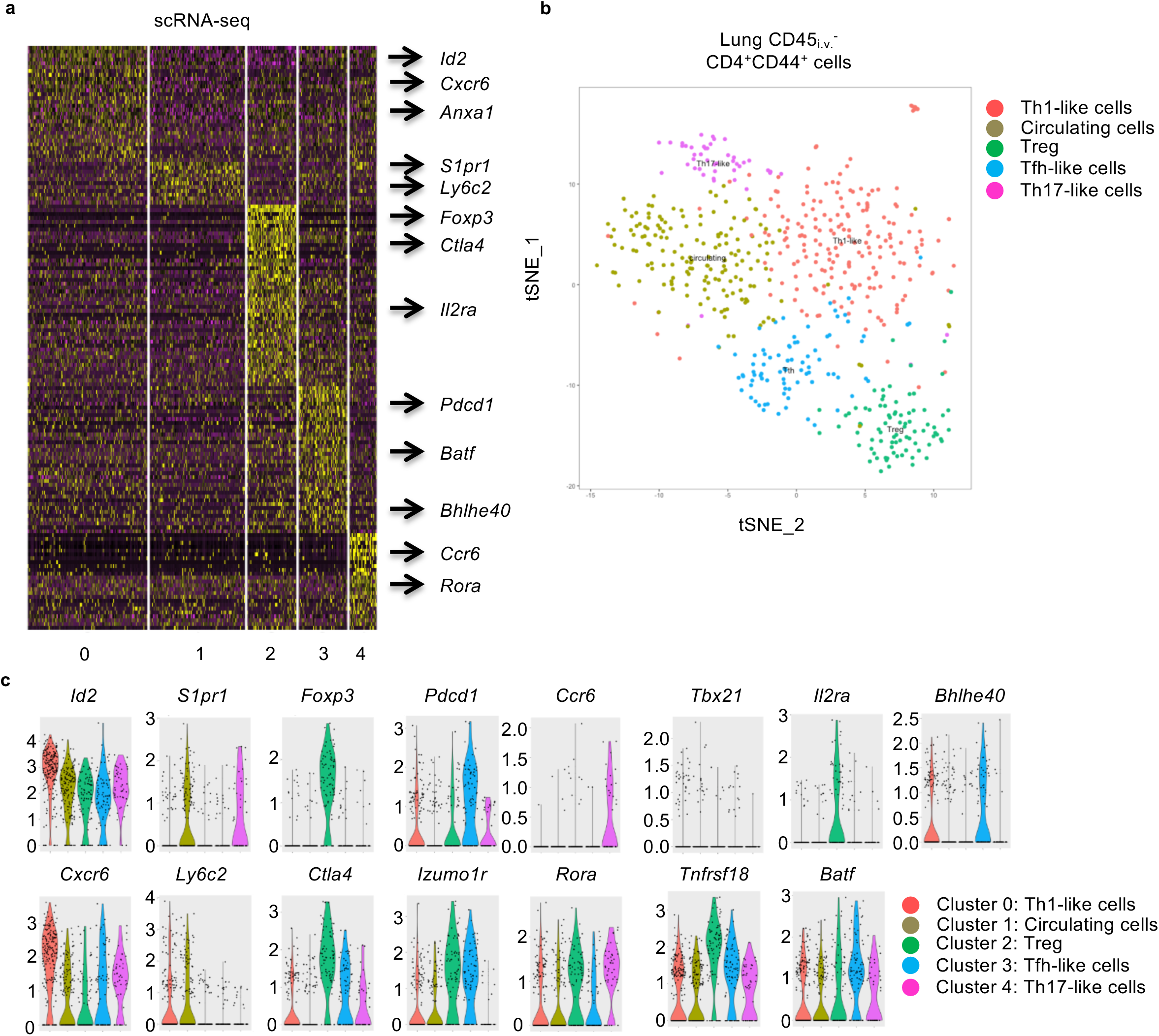
scRNA-seq identifies a population of lung T_H_ cells exhibit T_FH_ features. **a**, Heat map of signature genes following high-rank cluster analysis of scRNAseq data. **b**, 5 clusters of lung T helper cells were identified following scRNA seq analysis. **c**, Violin plot of representative T_H_ genes in each cluster.

**Extended Fig. 4.**
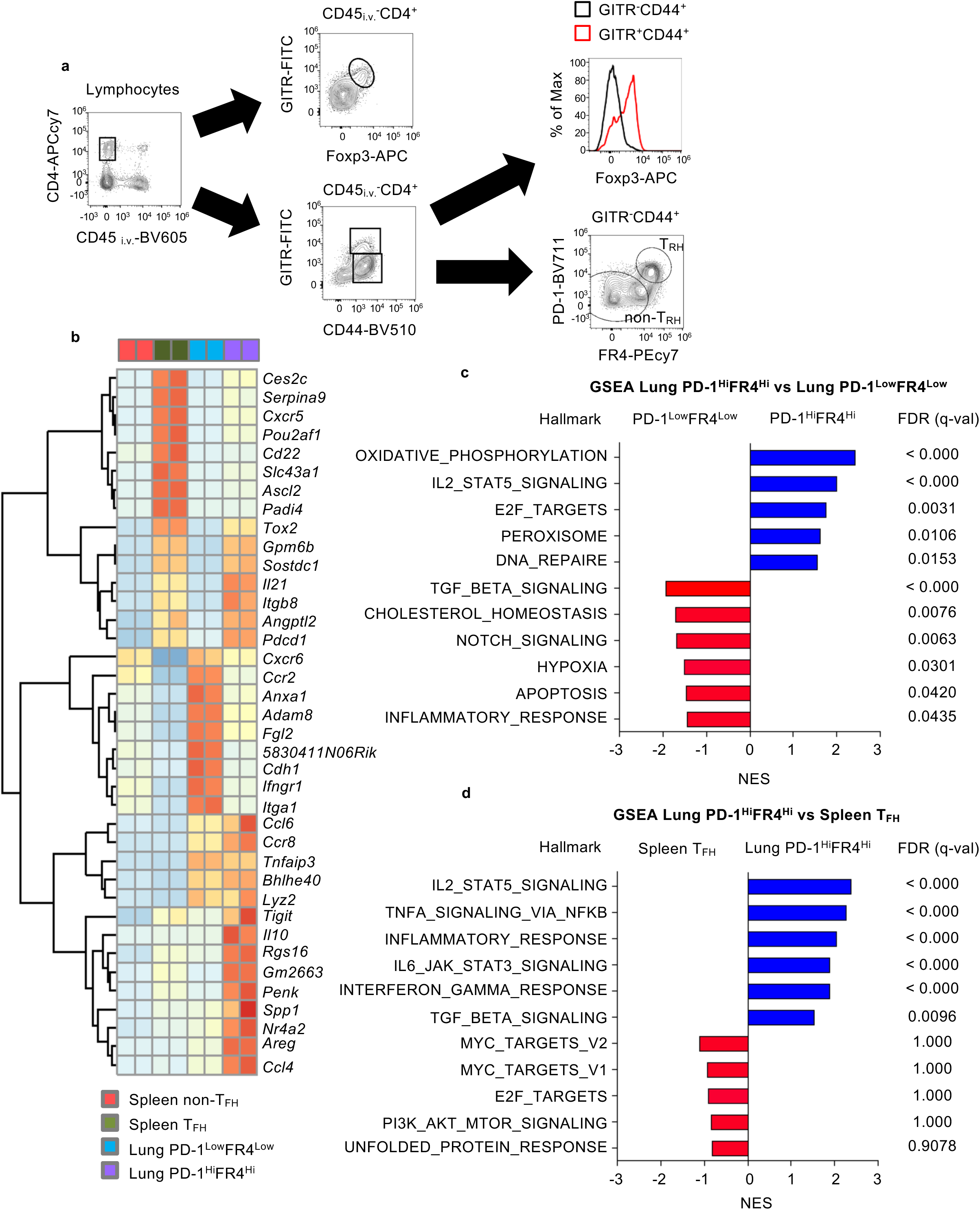
Total RNA-seq identified PD-1^Hi^FR4^Hi^ cells exhibits features of T_FH_ and T_RM_. **a**, Cell sorting strategy for lung PD-1^Hi^FR4^Hi^, PD-1^Low^FR4^Low^ population in the lungs. To exclude Foxp3^+^ T_reg_ cells, lung PD-1^Hi^FR4^Hi^, PD-1^Low^FR4^Low^ cells were sorted from CD45_i.v._^-^CD4^+^GITR^-^ CD44^Hi^ population. **b**, Cluster analysis of signature genes in lung PD-1^Hi^FR4^Hi^, lung PD-1^Low^FR4^Low^, splenic T_FH_ or splenic non-T_FH_ cells. **c**, GSEA of lung PD-1^Hi^FR4^Hi^ and lung PD-1^Low^FR4^Low^ cells. **d**, GSEA of lung PD-1^Hi^FR4^Hi^ and splenic T_FH_ cells.

**Extended Fig. 5.**
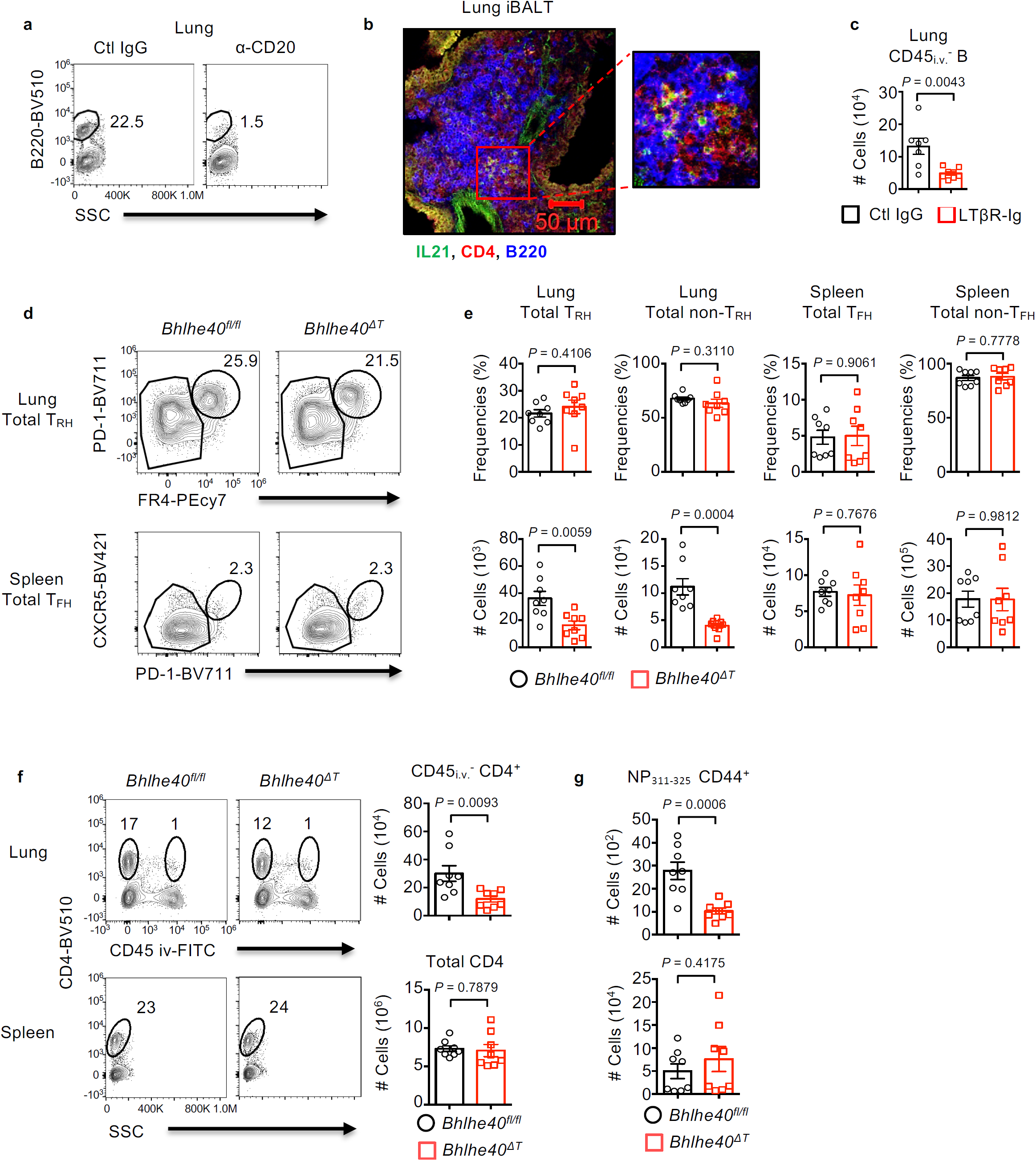
Impaired T_RH_ responses following B cell depletion, iBALT disruption or Bhlhe40 deficiency. **a-c**, WT (**a** and **c**) or IL-21-VFP reporter (**b**) mice were infected with PR8 and received with ctl IgG, α-CD20 or LTβR-Ig weekly in the present of FTY-720 (Experimental scheme in Fig 5. a.). **a**, The efficiency of B cell depletion in the lung. **b**, Representative confocal images of IL-21-expressing cells in iBALT. **c**, Cell numbers of lung tissue B cells in mice received with ctl-IgG or LTβR-Ig. **d-g**, *Bhlhe40*^*fl/fl*^ or *Bhlhe40*^*ΔT*^ mice were infected with PR8. Representative dot plot (**d**), percentages (top) or cell numbers (bottom) (**e**) of total lung T_RH_, non-T_RH_, splenic T_FH_ or splenic non-T_FH_ at 28 d.p.i. **f**, Representative dot plot or cell numbers of lung tissue or splenic CD4^+^ T cells at 28 d.p.i. **g**, Cell numbers of lung or splenic influenza-NP_311-325_ CD4^+^ T cell at 28 d.p.i. Data were pooled from two independent experiments (3-4 mice per group). *P* value were calculated by unpaired two-tailed Student’s t-test.

**Extended Fig. 6.**
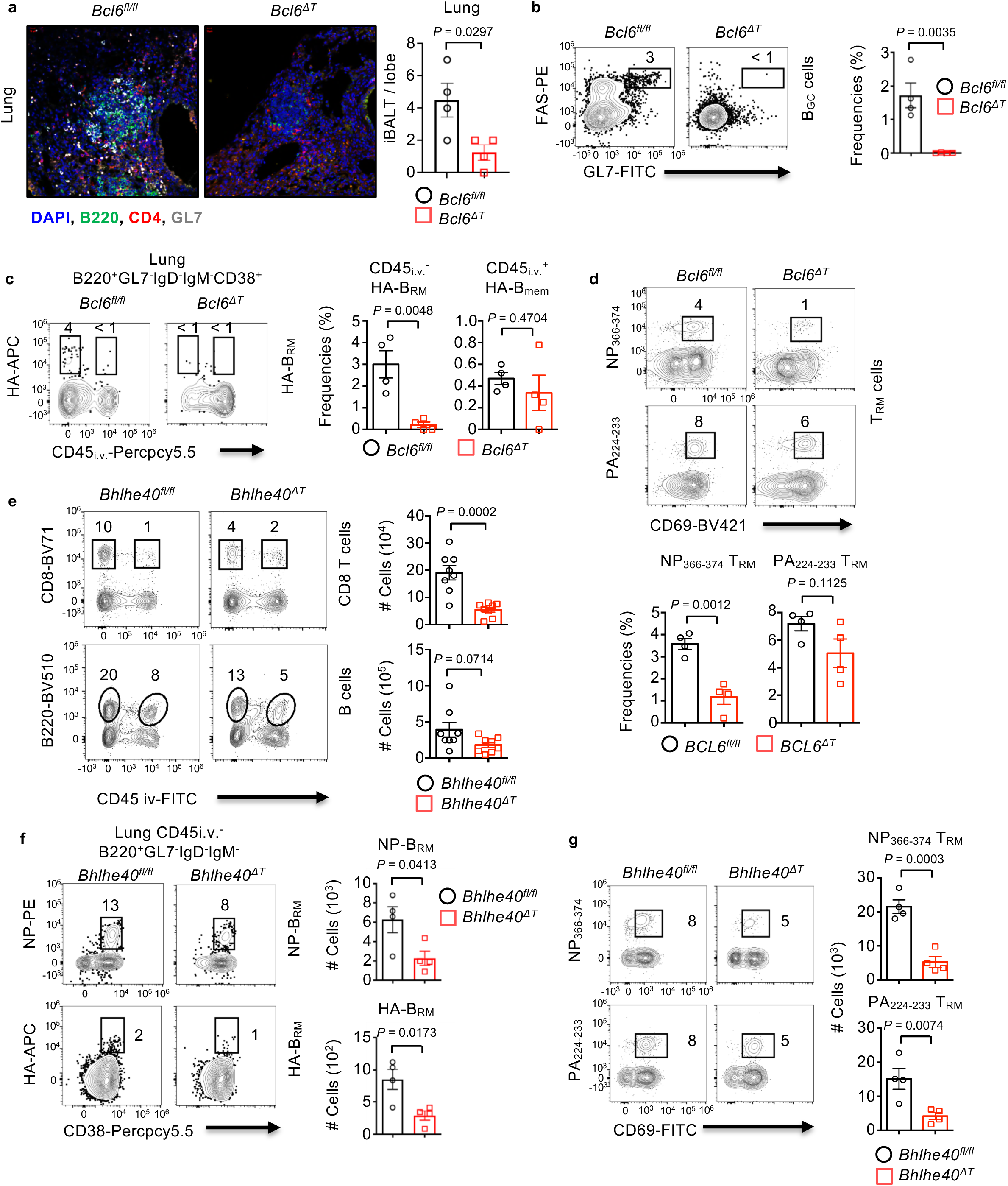
T cell-specific BCL6 or Bhlhe40 deficiency leads to impaired lung CD8^+^ memory and B cell immunity. **a-d**, *Bcl6*^*fl/fl*^ or *Bcl6*^*ΔT*^ mice were infected with PR8. **a**, Representative confocal images of lung iBALT at 28 d.p.i. Lung sections were stained with α-CD4 (red), α-B220 (green), α-GL7 (white) and DAPI (blue). Percentages of lung B_GC_ (**b**), lung tissue HA-specific B_RM_ or circulating HA-specific B_MEM_ (**c**) and CD8^+^CD69^+^ NP_366-374,_ or CD8^+^CD69^+^ PA_224-233_ T_RM_ (**d**). **e-g**, *Bhlhe40*^*fl/fl*^ or *Bhlhe40*^*ΔT*^ mice were infected with PR8. **e**, Representative dot plot or cell numbers of lung tissue CD8^+^ T (top) or B cells (bottom) at 28 d.p.i. **f**, Cell numbers of NP-specific B_RM_ or HA-specific B_RM_ cells. **g**, Cell numbers of CD8^+^CD69^+^ NP_366-374,_ or CD8^+^CD69^+^ PA_224-233_ T_RM._. In **a-d** and **f-g**, representative data were from at least two independent experiments (4-5 mice per group). In **e**, data were pooled from two independent experiments (4 mice per group). *P* values of all experiments were calculated by unpaired two-tailed Student’s t-test.

**Extended Fig. 7.**
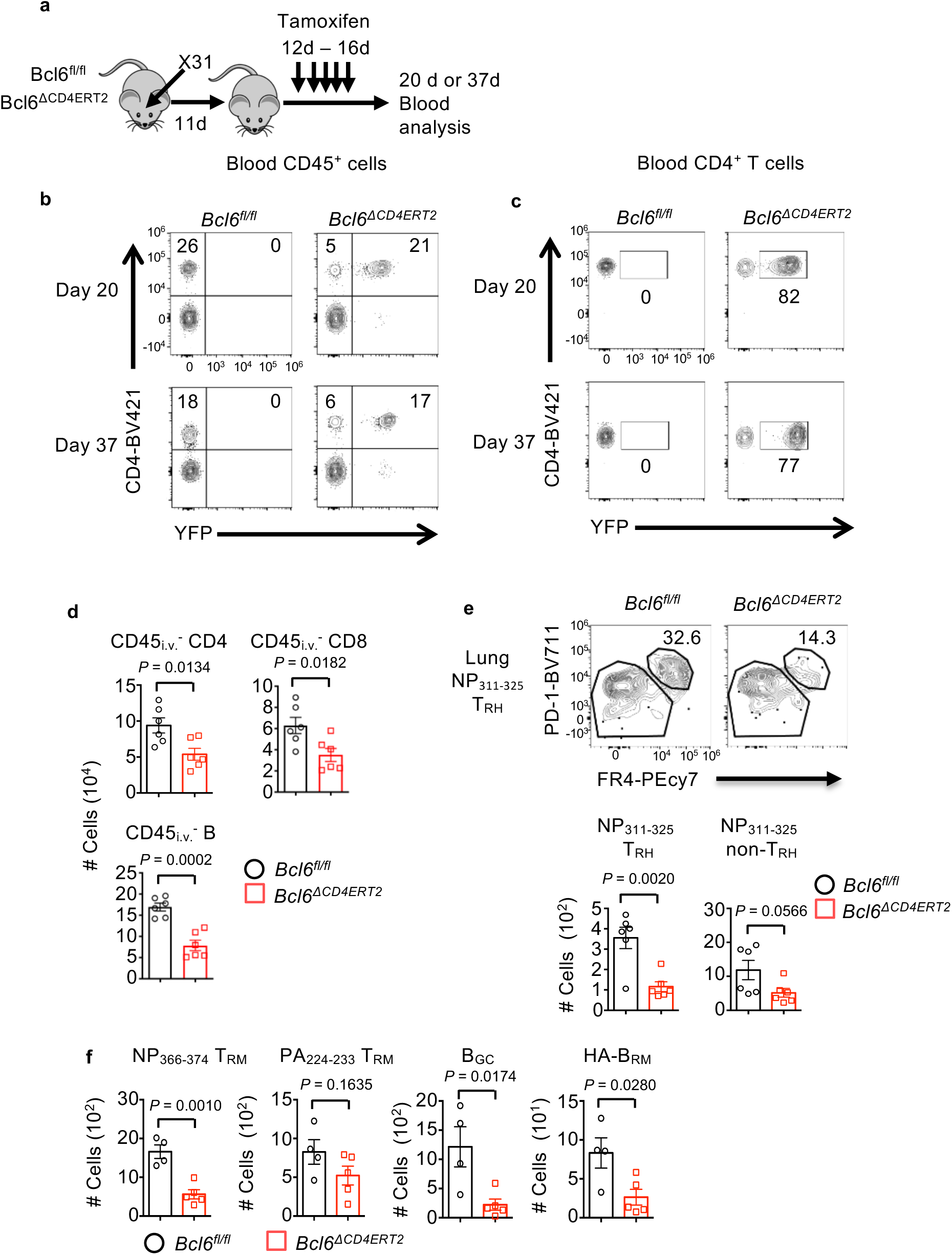
T_RH_ cells help local development of memory CD8^+^ and B cells. **a-c**, *Bcl6*^*fl/fl*^ (with ROSA26 LSL-YFP transgene) or *Bcl6*^*ΔCD4ERT2*^ (with ROSA26 LSL-YFP transgene) mice were infected with X31. Tamoxifen was administrated daily from 12 to 16 d.p.i. **a**, Experimental scheme of selective deletion of *Bcl6* in CD4^+^ T cells. **b, c**, YFP expression in blood CD45^+^ (**b**) or CD4^+^ T (**c**) cells following tamoxifen injection. **d, e**, *Bcl6*^*fl/fl*^ or *Bcl6*^*ΔCD4ERT2*^ mice were infected with PR8. Tamoxifen was administrated daily from 12-16 d.p.i. in the presence of daily FTY720 administration (11-34 d.p.i.). **d**, Cell numbers of CD45_i.v._^-^ CD4^+^ T, CD8^+^ T or B cells at 35 d.p.i. **e**, Representative dot plot (top) or cell numbers (bottom) of NP_311-_ 325 T_RH_ or non-T_RH_ cells at 35 d.p.i. **f**, Cell numbers of lung CD8^+^CD69^+^ NP_366-374_ or CD8^+^CD69^+^ PA_224-233_ T_RM_, B_GC_ or HA-specific B_RM_ cells were enumerated from X31-infected *Bcl6*^*fl/fl*^ and *Bcl6*^*ΔCD4ERT2*^ mice in the present of FTY720 (11-34 d.p.i.) at 35 d.p.i. In **a-c** and **f**, representative data were from at least two independent experiments (2-5 mice per group). In **d-e**, data were pooled from two independent experiments (3 mice per group). *P* values of all experiments were calculated by unpaired two-tailed Student’s t-test.

**Extended Fig. 8.**
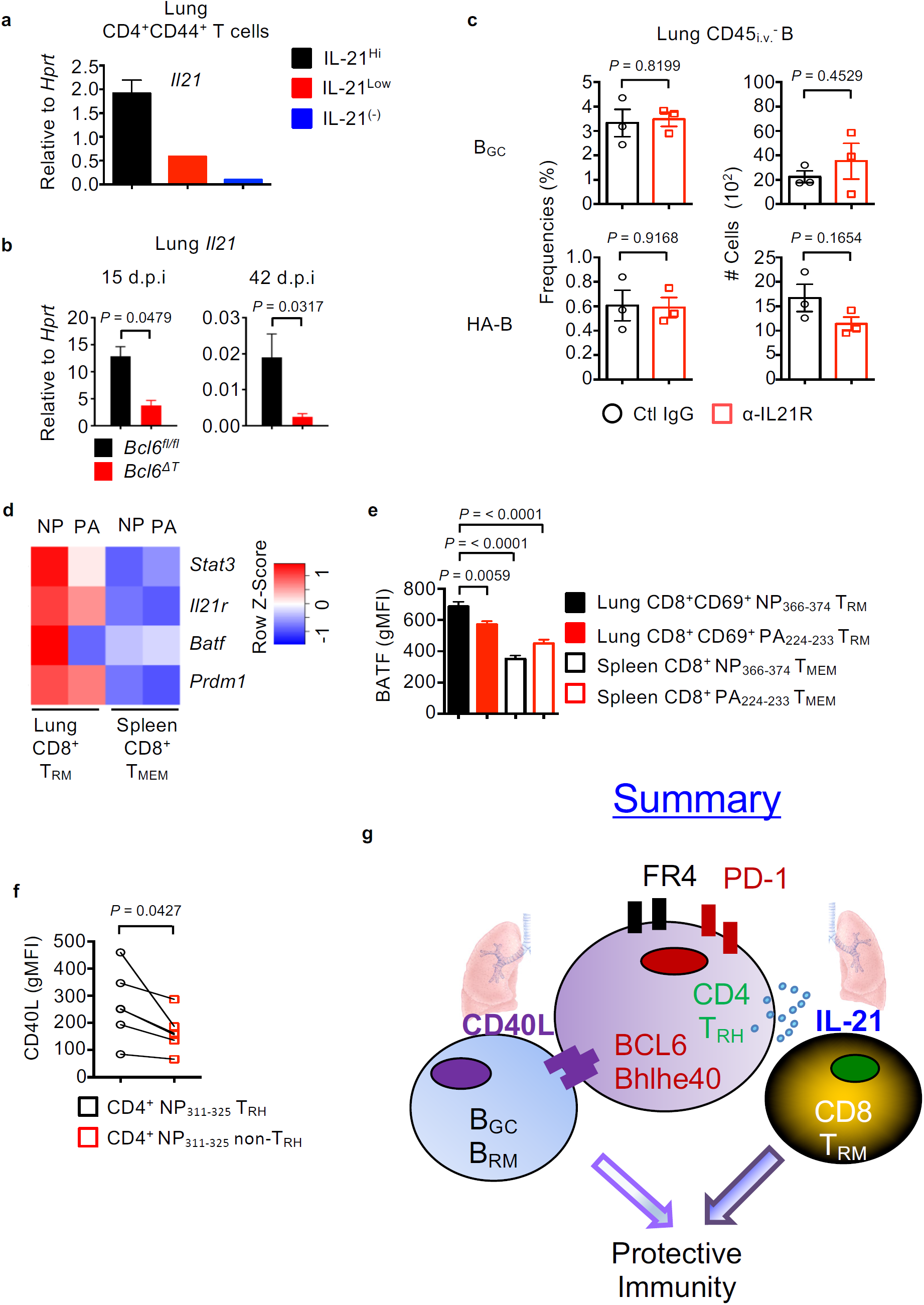
T_RH_ cells help CD8^+^ T and B cells through IL-21 or CD40L dependent mechanisms. **a**, Expression of *Il21* gene in sorted IL-21-VFP^Hi,^ IL-21-VFP^Low^ or IL-21-VFP^-^ CD4^+^ T cells from the lung tissue at 21 d.p.i. (pooled from 8 mice) **b**, Expression of *Il21* in the whole lung from *Bcl6*^*fl/fl*^ or *Bcl6*^*ΔT*^ mice at 15 or 42 d.p.i. **c**, Frequencies or cell numbers of lung B_GC_ or HA-specific B cells from PR8 infected WT mice treated with ctl-IgG or α-IL-21R (Intranasal route). **d**, Heat map of IL-21 signaling-related genes from sorted lung CD8^+^ NP_366-374_ or PA_224-233_ T_RM_ or splenic CD8^+^ NP_366-374_ or PA_224-233_ T_MEM_ determined by Nanostring at 38 d.p.i. as reported previously ^40^. **e**, Geometric M.F.I. of BATF expression in lung CD8^+^ NP_366-374,_ CD8^+^ PA_224-233_ T_RM_, splenic CD8^+^ NP_366-374_ or PA_224-233_ T_MEM_ cells from PR8 infected mice at 35 d.p.i. **f**, CD40L expression in the NP_311-325_ T_RH_ or non-T_RH_ cells following PMA/Ionomycin stimulation at 28 d.p.i. **g**, Schematics of the summary. In **b-c** and **e-f**, representative data were from at least two independent experiments (3-5 mice per group). *P* values were calculated by unpaired (b and c) or paired (f) two-tailed Student’s t-test. *P* value in e was analyzed by one-way ANOVA.

